# Chromosomal Mcm2-7 distribution is the primary driver of the genome replication program in species from yeast to humans

**DOI:** 10.1101/737742

**Authors:** Eric J. Foss, Smitha Sripathy, Tonibelle Gatbonton-Schwager, Hyunchang Kwak, Adam H. Thiesen, Uyen Lao, Antonio Bedalov

**Affiliations:** Clinical Research Division, Fred Hutchinson Cancer Research Center, Seattle, WA; Department of Medicine, Department of Biochemistry, University of Washington, Seattle WA

## Abstract

The spatio-temporal program of genome replication across eukaryotes is thought to be driven both by the uneven loading of pre-replication complexes (pre-RCs) across the genome at the onset of S-phase, and by differences in the timing of activation of these complexes during S-phase. To determine the degree to which distribution of pre-RC loading alone could account for chromosomal replication patterns, we mapped the binding sites of the Mcm2-7 helicase complex (MCM) in budding yeast, fission yeast, mouse and humans. We observed identical MCM double-hexamer footprints across the species, but notable differences in their distribution: In budding yeast, complexes were present in sharp peaks comprised largely of single double-hexamers; in fission yeast, corresponding peaks typically contained 4 to 8 double-hexamers, were more disperse, and showed a striking correlation with AT content. In mouse and humans, complexes were even more disperse, with a preference for regions of high GC content. Nonetheless, most fluctuations in replication timing in all four organisms could be accounted for by differences in chromosomal MCM distribution. This analysis also identified genomic regions whose replication timing was clearly not attributable to MCM density. The most notable was the inactive X-chromosome, which replicates late in S phase despite the fact that both MCM abundance and chromosomal distribution were comparable to those on the early replicating active X-chromosome. We conclude that, although certain genomic regions, most notably the inactive X-chromosome, are subject to post-licensing regulation, most differences in replication timing along the chromosome reflect uneven chromosomal distribution of stochastically firing pre-replication complexes.

## Introduction

Eukaryotes organize the replication of their genomes according to programs that ensure that certain regions of the genome complete replication earlier in S phase than others (Bleichert, Botchan, & Berger, 2017; Dileep & Gilbert, 2018; Fragkos, Ganier, Coulombe, & Mechali, 2015; Rhind & Gilbert, 2013; Riera et al., 2017). Differences in replication timing can have significant consequences, as late replication is associated with higher frequencies of mutation and genome rearrangement (Chen et al., 2010; Koren et al., 2012; Lang & Murray, 2011; Stamatoyannopoulos et al., 2009; Weber, Pink, & Hurst, 2012; Yehuda et al., 2018). The importance of replication timing is underscored by the observation that it is sometimes modulated according to the utility of the region being replicated: For example, some chromosomal regions containing developmentally regulated genes replicate early only during those developmental phases during which they are activated (Hiratani et al., 2010; Rivera-Mulia et al., 2015; Siefert, Georgescu, Wren, Koren, & Sansam, 2017). An even more striking example is the X chromosome in female mammals, where the active X chromosome (Xa) replicates early while the inactive X (Xi) replicates late (Gilbert, Muldal, Lajtha, & Rowley, 1962; Hansen, Canfield, Fjeld, & Gartler, 1996; Heard & Disteche, 2006).

The two main mechanisms for cellular control of replication timing are the regulation of (1) loading and (2) activation of the replicative helicase complex, also referred to as origin licensing and firing, respectively: Only sites where the replicative helicase has been loaded during G1 are capable of initiating replication during the subsequent S phase, and only a subset of these loaded helicases are actually activated (Bell & Labib, 2016; Fragkos et al., 2015). Furthermore, because there is a limiting supply of the proteins required for helicase activation, not all of the helicases that will be activated can be activated simultaneously, and firing factors are recycled to allow sequential firing of origins (Mantiero, Mackenzie, Donaldson, & Zegerman, 2011; Tanaka, Nakato, Katou, Shirahige, & Araki, 2011). Despite the biological importance of replication timing, the relative importance of helicase loading versus helicase activation in its control has not been established.

The replicative helicase consists of six highly related subunits, MCM2-7, whose N- and C-termini align to form a barrel-shaped structure with a central channel (Abid Ali et al., 2017; Noguchi et al., 2017). This hexamer is targeted to DNA at sites bound by the six-subunit Origin Recognition Complex (ORC), encoded by ORC1-6 (Bell & Labib, 2016). The sequences to which ORC binds differ between organisms: In the budding yeast *Saccharomyces cerevisiae*, it recognizes a relatively specific 30 base pair sequence that contains the 11 base pair ARS consensus sequence (ACS) (Eaton, Galani, Kang, Bell, & MacAlpine, 2010; Xu, Aparicio, Aparicio, & Tavare, 2006); in the fission yeast *Schizosaccharomyces pombe*, the Orc4 subunit contains AT hook domains, which target the complex to AT-rich sequences (Chuang & Kelly, 1999; Segurado, de Luis, & Antequera, 2003); and in metazoans, ORC does exhibit sequence preference, but instead recognizes some other feature of DNA or chromatin (Remus, Beall, & Botchan, 2004; Vashee et al., 2003). Regardless of these differences in the sites that ORC selects, the actual mechanism of loading of the helicase complex is the same in these organisms: ORC, Cdc6 and Cdt1 sequentially load two helicase complexes, using the energy of ATP to open each ring (Bell & Kaguni, 2013). This results in a double MCM hexamer encircling the DNA, oriented with the N-termini facing inward and the DNA entering and leaving through the channels at the C-termini. Activation of the loaded helicases occurs in response to the rising activity of two kinase complexes, DDK and CDK, and results in the two hexamer rings locked shut with single strands of DNA of opposite polarity going through the two central channels (Labib, 2010).

Here we identify, at nucleotide resolution, all sites at which replicative helicases have been loaded in budding yeast, fission yeast, mouse and humans. By comparing the distribution of these sites with the patterns of replication in the corresponding organisms, we found that, with a few notable exceptions, the pattern of replication timing in S phase is largely predictable simply on the basis of the pattern of helicase loading in the preceding G1. This suggests that control of replication timing is simpler than has been widely presumed, with most of the regulation occurring at the level of helicase loading, and the subsequent firing of origins being largely stochastic.

## Results and Discussion

### Mcm-ChEC identifies single MCM double-hexamer footprints adjacent to ACS at *S. cerevisiae* origins

The Mcm2-7 complex encircles approximately 60 base pairs (bp) of DNA as a double-hexamer (DH) whose monomer components are juxtaposed head-to-head at their N-termini with C-termini facing away from each other (Abid Ali et al., 2017). To exploit this arrangement of Mcm2-7 hexamers as a way to identify their exact loading sites *in vivo*, we tagged Mcm2, Mcm4 and Mcm6 subunits at their C-termini in *Saccharomyces cerevisiae* with micrococcal nuclease (MNase) (Schmid, Durussel, & Laemmli, 2004; Zentner, Kasinathan, Xin, Rohs, & Henikoff, 2015), permeabilized G1-arrested cells, activated the MNase by adding calcium, and prepared libraries for paired end sequencing using total extracted DNA without any size fractionation (Foss et al., 2019). Strains with tagged Mcm proteins exhibited growth rates comparable to those in wild type, indicating that the presence of the tag did not perturb protein function (Supplementary data Fig. 1). Because PCR involved in library preparation preferentially amplifies short fragments, which in turn are generated by MNase activity, we predicted that resulting libraries would reflect the sites at which Mcm2-7 DHs had been loaded; furthermore, these sites should be congruent for the three different libraries and correspond, at least partially, to known replication origins. Consistent with this expectation, our sequencing results for libraries prepared from the three different tagged subunits were focused in sharp peaks that coincided with each other (Fig 1A) and with known replication origins and replication initiation sites (Fig. 2A, left middle panel). Furthermore, fragment size distribution for all three libraries peaked in the 50-62 bp size range, consistent with the 60-62 bp protected by Mcm2-7 DHs observed in cryo-electron microscopy studies (Abid Ali et al., 2017; Noguchi et al., 2017) (Supplementary data Fig. 2). We interpret fragment peaks in the 150-200 bp size range as reflections of cleavage by the MNase-tagged protein between flanking nucleosomes and note that such fragments have been observed for other MNase tagged proteins, including transcription factors (Foss et al., 2019). In this report, we visualize individual Mcm2-7 footprints as heat maps, with fragment size plotted according to genomic location, and read depths represented by color intensity (Foss et al., 2019). A typical example is shown in Fig. 1B: The Mcm2 footprint at ARS1103 was composed of reads in the 50-70 bp size range located immediately adjacent to the ARS consensus sequence (ACS) bound by the ORC, which loads the MCM complex. We found similar Mcm2 footprints in each of the 343 origins listed in the Saccharomyces Genome Database (SGD), and in all 187 instances in which the origin had a defined ACS, the footprint was located immediately adjacent to the ACS. All *S. cerevisiae* origins are shown in Supplementary Data Fig. 3. Essentially identical footprints were generated by MNase-tagged Mcm4 and Mcm6 (Fig 1B), confirming that footprints reflect Mcm2-7 binding. Furthermore, G1-specific and Cdc6-dependent footprints obtained by sequencing DNA fragments from nuclei treated with exogenous MNase (MNase-seq) coincided with these Mcm-MNase generated signals (Foss et al., 2019). The prominent Mcm2 signal at the highly efficient ARS1103 origin (Fig. 2A, left middle panel red arrow) suggests that at least one Mcm2-7 DH is loaded at this site in many cells, and the virtual absence of an Mcm2-7 signal in the surrounding 1.5 kb of DNA rules out the presence of multiple Mcm2-7 complexes at this origin. We conclude that replication origins do not require multiple MCM complexes in order to be highly efficient, and that variations in the MCM signal that we observe across the genome predominantly reflect variation in the fraction of cells in which a single complex has been loaded (all *S. cerevisiae* origins are shown in Supplementary Data Fig. 3). Low abundance Mcm2-7 signals found outside known replication origins also feature footprints of a single DH and flanking nucleosomes (Supplementary Data Fig. 4), suggesting that low levels of DNA replication may initiate from a plethora of genomic sites.

**Figure 1.**
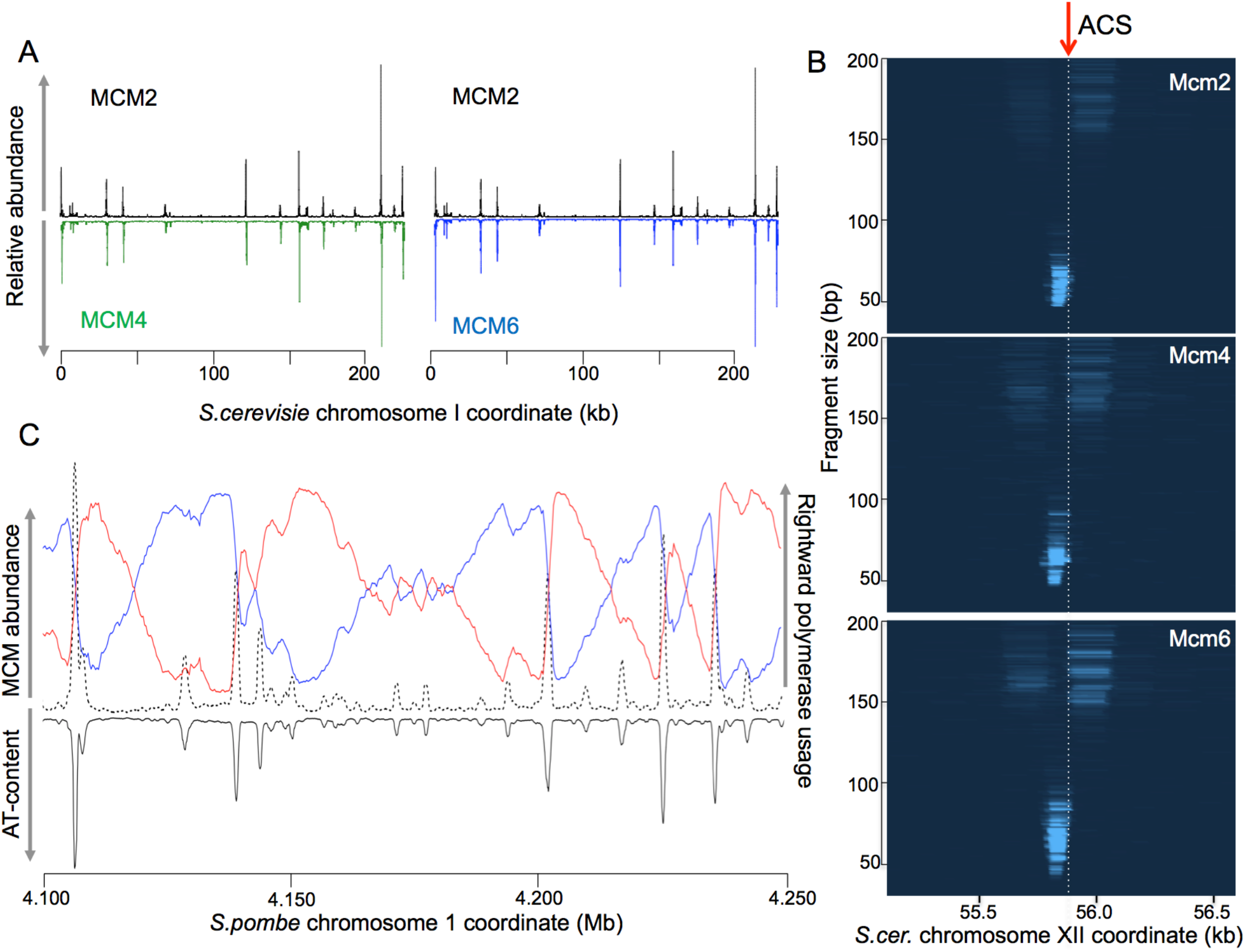
Tagging of MCM subunits with MNase in yeast reveals MCM helicase binding sites. (A) Plot of read depths across chromosome I showing that binding sites for Mcm2 (black, top panels), Mcm4 (green, bottom left) and Mcm6 (blue, bottom right) are largely congruent. (B) MCM subunit binding sites, as characterized by chromosomal location on the X-axis, fragment size on the Y-axis and read depth, based on color intensity, and illustrated by images reminiscent of agarose gels. All three subunits generate similar footprints. Shown is ARS1103 on chr. XI. (C) Mcm2-ChEC (dotted black line), polymerase usage (blue and red lines; data from Daigaku *et al.* (Daigaku et al., 2015)) and AT content (solid black line) for 150 kb segment of *S. pombe* chr. II. Red and blue lines show fraction of newly synthesized Watson strand that was generated by pol epsilon and pol delta, respectively.

**Figure 2.**
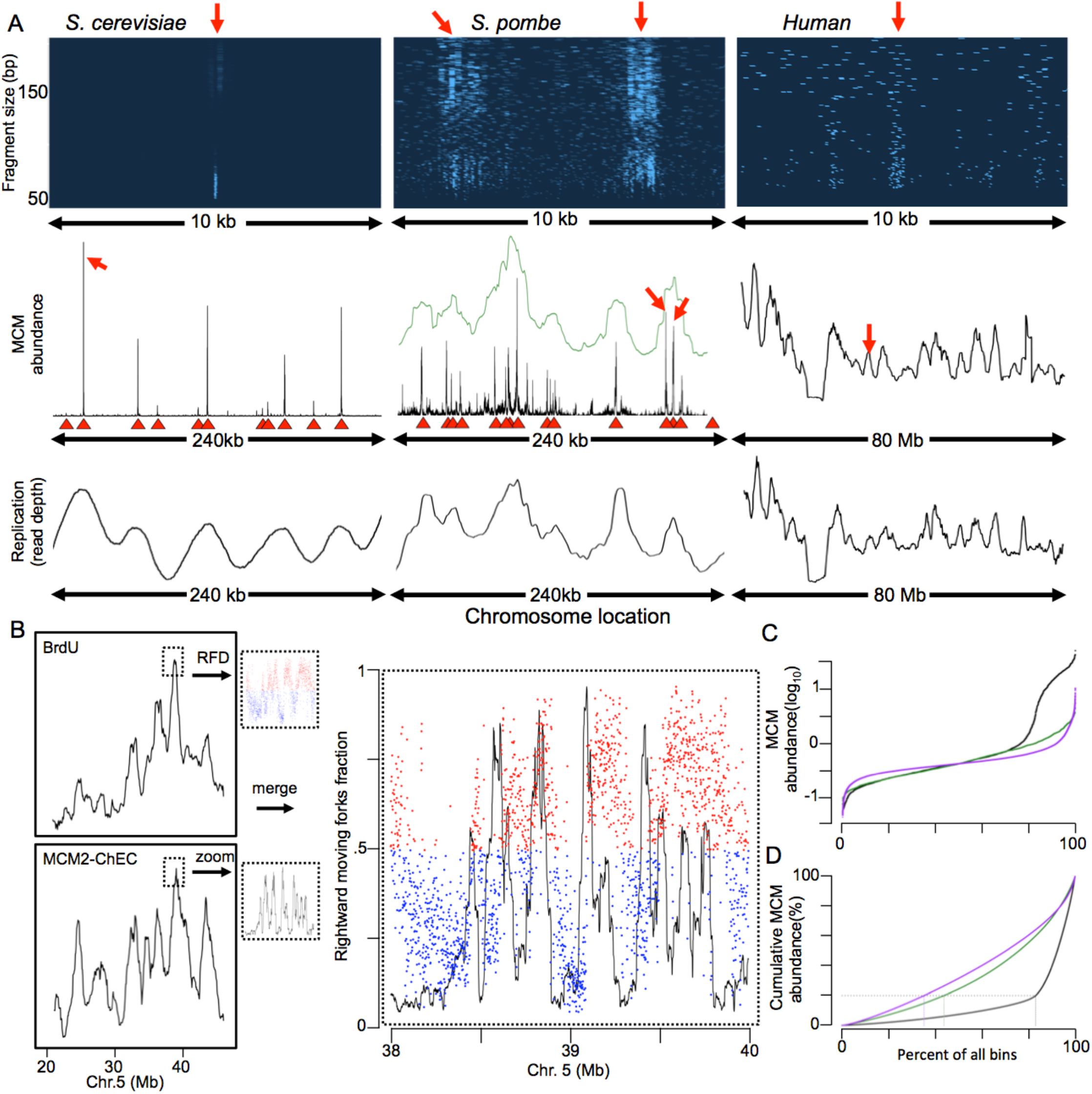
MCM distribution and replication in yeasts and human. (A) Left 3 panels show budding yeast (chr. XI), middle three panels show fission yeast (chr. II) and right 3 panels show HeLa cells (chr. 18). For Mcm2-ChEC experiments (top 2 rows), budding yeast were arrested for 2 hours with alpha factor, log phase fission yeast were transferred to medium containing 15 mM HU for 3 hours, and HeLa cells were arrested with contact inhibition. Top row: Mcm2 binding visualized by plotting chromosomal location on the x axis, fragment size on the y axis, and read depth indicated by color intensity. The budding yeast image corresponds to ARS1103 and the fission yeast to II-448. Middle row: Mcm2-ChEC read depths plotted across 240 kb (both yeasts) or 80 Mb (HeLa). Green line in the middle plot (*S. pombe*) shows Mcm2-ChEC read depths divided into 1 kb bins and smoothed with a 15 bin sliding window. Red triangles in both yeasts indicate known origins of replication, as listed in oridb.org. Red arrows indicate regions of correspondence between the top and middle rows. Bottom row: Replication as determined by S to G1 ratio of flow sorted cells for budding and fission yeast, and by BrdU incorporation for HeLa. (B) Replication, as determined by BrdU incorporation (Davis et al., 2018), (upper left panel) and MCM binding (lower left panel) is shown for HeLa cells. Right panel shows blow-up of indicated area with fraction of forks involved in synthesizing the Watson strand that are moving to the right (red and blue dots are used to emphasize direction of majority of synthesis; data from Petryk *et al.* (Petryk et al., 2016)) and MCM binding (black). Blue to red transitions denote the initiation zones, which coincide with local MCM maxima. (C) MCM distribution expressed as log_10_ fraction of median value for 5 kb-binned Mcm2-ChEC signal. *S. cerevisiae, S. pombe*, and HeLa are shown in black, green and purple, respectively. (D) Cumulative distribution of 5 kb-binned Mcm2-ChEC signal in *S. cerevisiae* (black), *S. pombe* (green), and HeLa (purple).

### AT-content dictates chromosomal MCM distribution and replication initiation in *S. pombe*

We used the same approach to identify Mcm2-7 localization in the fission yeast *Schizosaccharomyces pombe*. As in budding yeast, the distribution of fragment lengths was bimodal, with the major peak consistent with the size protected by MCM complexes (Supplementary data Fig. 5). Also like in budding yeast, the Mcm2 peaks largely coincided with previously identified origins of replication (Fig. 2A, central panel). Furthermore, Fig. 1C demonstrates a striking preference of MCM binding for AT-rich sequences, which is consistent with observations that (1) replication in *S. pombe* initiates predominantly from AT-rich sequences (Segurado et al., 2003), and (2) fission yeast ORC binds preferentially to AT-rich sequences due to AT hook domains in the Orc4 subunit (Chuang & Kelly, 1999). However, two features of Mcm2-7 distribution in fission yeast differed markedly from budding yeast: First, unlike *S. cerevisiae*, where we found a single MCM DH at most origins, we found 4-12 adjacent DH distributed over 500-1500 bp at most known *S. pomb*e origins (Fig 2A, top middle panel; all *S. pombe* origins are shown in Supplementary data Fig 6). Second, we found significantly more minor MCM peaks between origins (center versus left panel in middle row of Fig. 2A). To determine whether the observed pattern of MCM distribution matches the pattern of replication initiation, we aligned MCM distribution with previously published *S.pombe* DNA polymerase usage sequence analysis (Pu-seq) datasets (Daigaku et al., 2015). Pu-seq identifies replication initiation sites, at high resolution, as DNA regions with sharp transition in Pol-epsilon and Pol-delta usage. As can be seen in Fig. 1C, we observed a striking co-localization among MCM and AT-abundance peaks with the polymerase usage transition sites. This is evident both for major and minor MCM peaks. We conclude that the AT-content dictates chromosomal MCM distribution, which in turn drives the replication initiation pattern in *S. pombe*.

### MCM distribution drives replication initiation pattern of human chromosomes

We next determined the distribution of Mcm2-7 complexes in human cells using MNase-seq. To do so, we expressed human Mcm2 tagged with MNase in HeLa cells using a lentiviral vector, choosing an expression level that did not exceed physiological levels (Supplementary data Fig. 7). As seen in yeast, the distribution of sequenced fragment sizes was bimodal, with the major peak composed of 50-100 bp fragments (Supplementary data Fig. 8). We observed individual Mcm2-7 footprints similar to those in both yeast species (Fig. 2A, upper right), indicating that the basic architecture of the complex is conserved. However, despite the similar appearance of individual footprints, the MCM signal in HeLa cells was more disperse than in yeast. After partitioning the genome into 5 kb bins and ordering bins based on MCM abundance, we observed a 9-fold difference in median MCM abundance between the top and bottom 5% of bins in human cells, compared to 11-fold and 94-fold difference in *S. pombe* and *S. cerevisiae* (Fig. 2C), respectively. The more disperse distribution of DHs in HeLa cells was also evident by examination of cumulative MCM distribution, with the top 65% of bins constituting 80% of the total Mcm2-7 signal, compared to 56% and only 17% in fission and budding yeast, respectively (Fig. 2D). These results demonstrate that while the basic architecture of binding of MCM DHs is preserved across species, their distribution becomes progressively more dispersed from budding yeast to fission yeast to mammals.

We next asked whether chromosomal regions with higher MCM density were correlated with earlier replication, using published data to assess replication timing. In a previous study (Chen et al., 2010), asynchronously growing HeLa cells were given a BrdU pulse and then flow-sorted based on DNA content; DNA was then immunoprecipitated with anti-BrdU antibodies and subjected to deep sequencing. As is clear from our analysis of these data (Fig. 2A, bottom row), although human chromosomes are approximately 100 times as long as those in yeast, the number of early- and late-replicating regions on a single chromosome is similar in both organisms, varying from 5 to 20 depending on the chromosome size. Chromosomal distribution of MCM complexes in HeLa cells showed a striking correlation with the chromosomal replication pattern, as shown in the middle and lower right panels of figure 2A, for the entire 80 Mb of chromosome 18. Previous studies have shown that at a finer scale, replication within a single early- or late-replicating zone is not uniform; for example, studies using Okazaki fragment sequencing and BrdU incorporation have shown that replication initiation within these zones is concentrated in approximately 100 kb sub-zones (Petryk et al., 2016). Just as we saw increased MCM density in multi-megabase early replicating chromosomal regions (Fig 2A right panels), we observed local MCM maxima that coincided strikingly with initiation zones within a single broad region (Fig 2B and Supplementary data Fig. 9). We therefore conclude that in human cells, just like in *S. pombe*, chromosomal MCM density in G1 cells predicts the replication program in the ensuing S-phase at both gross and fine scales.

### Replication modeling in *S. cerevisiae* identifies repetitive DNA as well-licensed but late replicating chromosomal regions

While the chromosomal MCM distribution in both *S. pombe* and human cells clearly recapitulates the corresponding replication pattern (Fig. 2A middle and left panels), the discretely focused distribution of MCM binding in *S. cerevisiae* makes the link between chromosomal MCM density and chromosomal replication less apparent. To determine whether chromosomal MCM density in G1 is sufficient to drive replication timing pattern in budding yeast, we modeled Mcm2-7 binding and DNA replication *in silico*, assuming that in each cell DNA replication initiates at random from sites bound by Mcm2-7 in that cell. This model was remarkably accurate in predicting replication pattern and was robust to changes in additional assumptions about how limitations of origin firing factors restrict the number of origins that can fire simultaneously and how freely those limiting firing factors diffuse (Supplementary data Fig. 10; algorithm described in Methods). However, this analysis also identified chromosomal regions whose replication was not congruent with MCM density. Most notable were repetitive regions, including the *rDNA* and *CUP1* arrays as well as regions near chromosome ends, which replicated later than expected based on our MCM density-driven replication simulations (Supplementary data Fig. 10). Several of these well-licensed but late replicating regions in yeast, such as rDNA and telomeres, are subject to regional, gene-independent transcriptional silencing and, as such, represent the equivalent of metazoan heterochromatin.

### Identical MCM chromosomal abundance, distribution and replication order along early replicating active and late replicating inactive X-chromosome

Given that the largest piece of heterochromatin in mammals is the late replicating, inactive X-chromosome (Xi), we asked whether reduced Mcm2-7 loading or delayed origin activation underlies late replication of that chromosome. To do so, we employed a clonal *M. musculus*/*M. spretus* hybrid cell line (Berletch et al., 2015) to compare allele-specific replication dynamics and Mcm2-7 loading between active and inactive X-chromosomes. In these cells, the active X chromosome is derived exclusively from the *M. spretus* parent. Given that the *musculus* and *spretus* genomes contain a single nucleotide polymorphism (SNP) every 50-100 bp, both replication and MCM binding on the two parental chromosomes can be followed independently. We observed only minor allelic differences in replication between autosomes, as previously reported using a similar assay (Dileep & Gilbert, 2018). Replication timing of the active X-chromosome was similar to that of autosomes, while replication of the inactive musculus X-chromosome was markedly delayed (Fig. 3A, upper right). However, in contrast to observed differences in replication timing, we detected no difference in Mcm2-7 abundance on the inactive and active X chromosomes (Fig. 3A, lower right; ratio of spretus to musculus reads was 1.00 and 1.03 for chromosomes X and 16, respectively). This result demonstrates that delayed replication of the inactive X-chromosome is not a consequence of reduced origin licensing but, rather, of delayed origin activation.

**Figure 3.**
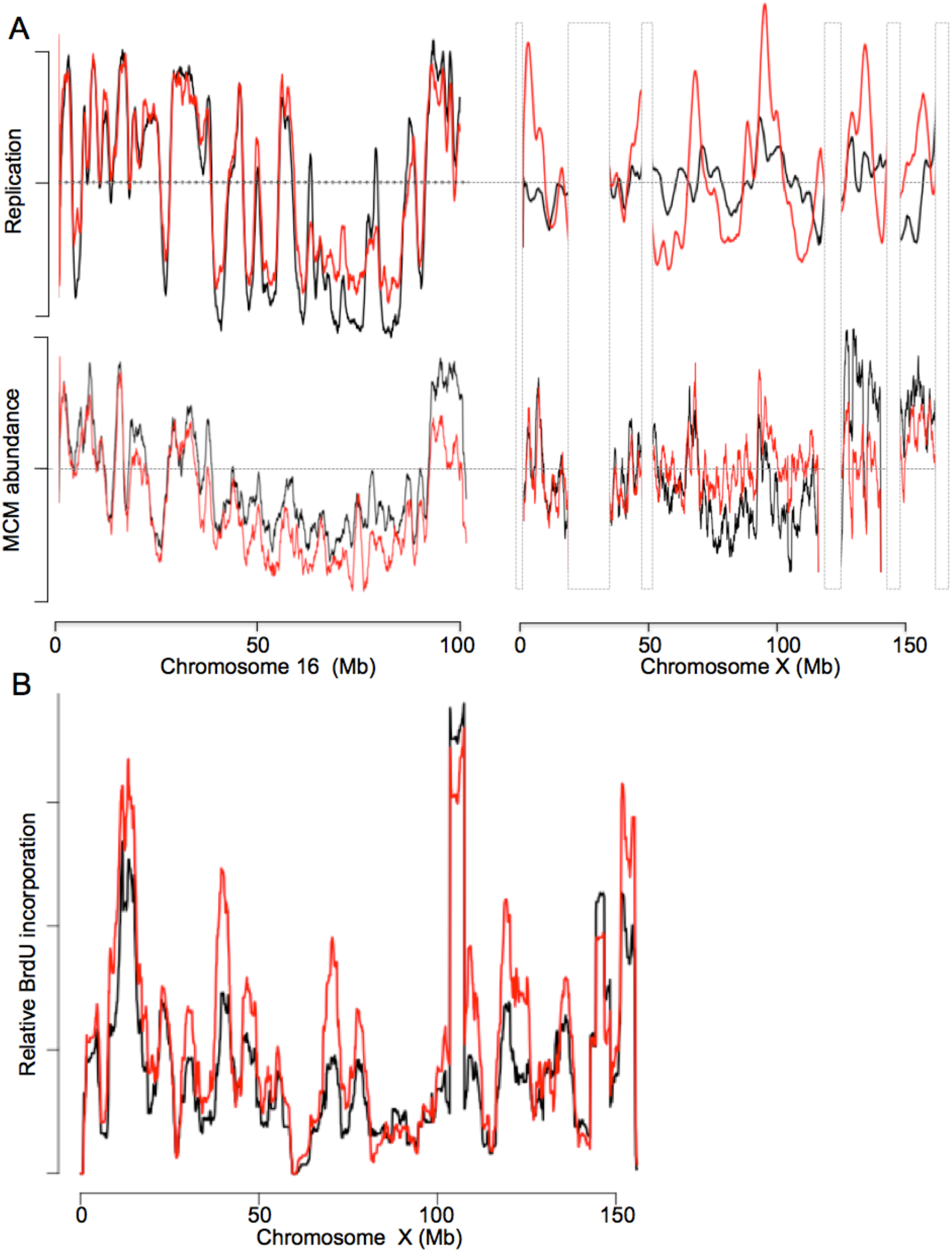
MCM distribution and replication on the X chromosome. (A) Replication (upper panel) and MCM binding (lower panel) in the mouse hybrid Patski cells for chr16 (left) and chrX (right). *M. musculus* and *M. spretus* parental chromosomes are shown in black and red, respectively. Replication is plotted as fraction of median read depth, with larger fluctuations from the median reflecting more active replication. Scale for MCM2 binding is arbitrary. (B) Fluctuation in BrdU incorporation across two copies of chr. X in human female cell line GM12878. Relative levels are normalized for the two chromosomes (data from ENCODE consortium (Davis et al., 2018)).

Despite differences in replication timing between active and inactive X chromosomes, the relative order in which different regions within each chromosome complete replication is preserved, with earlier replicating regions corresponding to regions of higher MCM density (Fig. 3A, right). This suggests that MCM density along the Xi determines the relative order of replication along the inactive X, as it does on the active X. Because this orderly replication pattern for the mouse Xi contrasts with the random initiation pattern others have reported for the human Xi (Koren & McCarroll, 2014), we re-examined Xi replication kinetics of in the human female lymphoblastoid cell line GM12878. This line is ideally suited for this purpose because (1) high-quality haplotype-resolved genome sequences are available (Eberle et al., 2017); (2) the active X chromosome is derived almost exclusively from one of the two parents (Kucera et al., 2011); and (3) high resolution replication profiles of these cells have been obtained by repli-seq (Davis et al., 2018). In the repli-seq study, BrdU incorporation was measured at several time points during S-phase, assuring similar resolution of replication kinetics within early and late S-phase. Our analysis of the repli-seq dataset showed that, as in the mouse, although the Xi replicates later than the active X, the relative order of replication along active and inactive chromosomes is comparable (Fig 3B). Given that transcription is largely abolished on the Xi, this observation suggests that the spatial pattern of both MCM loading and replication order on the Xi are independent of transcription. Furthermore, this result demonstrates that heterochromatin is not an impediment for MCM loading but rather for activation of licensed origins.

### GC-content, MCM distribution and replication initiation pattern of mammalian chromosomes

What determines chromosomal MCM distribution in mammals and how do these determinants differ from those that guide MCM distribution in lower eukaryotes? The MCM distribution in both budding and fission yeast is governed, in large part, by the binding preference of ORCs for specific sequences, namely, ACS in budding yeast (Hoggard, Shor, Muller, Nieduszynski, & Fox, 2013), and AT-rich sequences in fission yeast (Segurado et al., 2003), and this preference is clearly reflected in our results, as shown in Fig. 1B and Supplementary data Fig. 3 (*S. cerevisiae*) and Fig. 1C (*S. pombe*). In mammals, it is less clear whether MCM loading is guided by DNA sequence. On the one hand, the mammalian ORC reportedly does not exhibit sequence preferences but instead recognizes certain chromatin or DNA topological features (Remus et al., 2004; Vashee et al., 2003). On the other, several studies observed a bias toward GC-rich sequences in terms of both ORC localization and replication initiation (Cayrou et al., 2011; Dellino et al., 2013; Langley, Graf, Smith, & Krude, 2016; Miotto, Ji, & Struhl, 2016). Our observation that active and inactive X chromosomes, which differ vastly in chromatin organization but are almost identical in sequence, load MCM essentially identically favors the idea that DNA sequence or sequence-driven topological characteristics guide MCM distribution in mammals. Indeed, we found that GC content is highly correlated with MCM abundance in HeLa cells (r = 0.59), and with both ORC localization (r = 0.70) and replication initiation (r = 0.83), consistent with prior reports (Cayrou et al., 2011; Dellino et al., 2013; Langley et al., 2016; Miotto et al., 2016). (We note that the opposing trends in sequence preference that we observe here for MCM binding in *S. pombe* and mammals rule out the possibility that our results reflect the sequence preference of MNase.) To further assess the role of primary sequence as a determinant of MCM binding, ORC localization and genome replication, we analyzed pairwise correlations between (1) MCM binding; (2) ORC abundance (Dellino et al., 2013); (3) DNA replication, based on BrdU incorporation (Davis et al., 2018); (4) GC content; (5) the presence of single-stranded DNA (ssDNA), as measured by a KMnO_4_-based assay (Kouzine et al., 2017); (6) tendency to form G4 quadruplexes, based on the “G4 hunter” algorithm (Bedrat, Lacroix, & Mergny, 2016); (7) DNase-hypersensitivity (Davis et al., 2018); and (8) histone modifications associated with ORC binding and replication initiation (Davis et al., 2018) (see each pairwise r value in Supplementary Data Table 1). MCM binding was positively correlated with all of these metrics, with r values ranging from 0.46 to 0.91 (Fig 4A). Fig 4B shows, at a 50 kb scale, MCM co-localization with ORC and with features previously associated with ORC enrichment, including DNAse I hypersensitivity and histone modifications that mark active chromatin, including H3K4me3, H3K4me2, and H3K4me1, H3K27ac and H3K9ac (Miotto et al., 2016). Principal component analysis (PCA), using two components that collectively explain 86% of the variance (Supplementary data Fig. 11), revealed tight co-variation of MCM, replication initiation and histone H3K4Me3 H3K4Me2 marks (Fig. 4C). These findings are consistent with the idea that MCM is a primary driver of replication and that both MCM localization and replication initiation are associated with promoters and/or the 5’ ends of transcription units (Fig. 4B) which are enriched for H3K4Me3 and H3K4Me2. PCA also revealed striking co-localization of ORC and ssDNA. This association was also evident in the hierarchical clustering analysis in which ORC, GC content, single stranded DNA and propensity for G4 formation formed a major sub-cluster (Fig 4A). High GC content is associated with a variety of non-B DNA structures in which single-stranded DNA is extruded (Bartholdy, Mukhopadhyay, Lajugie, Aladjem, & Bouhassira, 2015). Even though structural studies of yeast and human ORCs show that the ORC encircles duplex DNA (N. Li et al., 2018; Tocilj et al., 2017), ORC in both yeast and humans binds ssDNA, which stimulates its ATPase activity (Hoshina et al., 2013; Lee, Makhov, Klemm, Griffith, & Bell, 2000), and may function in the initial loading of the ORC onto DNA. One specific non-B DNA form associated with DNA of high GC content is the G-quadruplex; our observed correlation of ORC binding with G-quadruplex-prone sequences (r = 0.83) suggests that G-quadruplexes may be one of the DNA structures associated with high GC content that is attractive to the ORC. These findings underscore the importance of chromosomal DNA sequence composition to MCM distribution and chromosomal replication pattern from yeast to humans.

**Figure 4.**
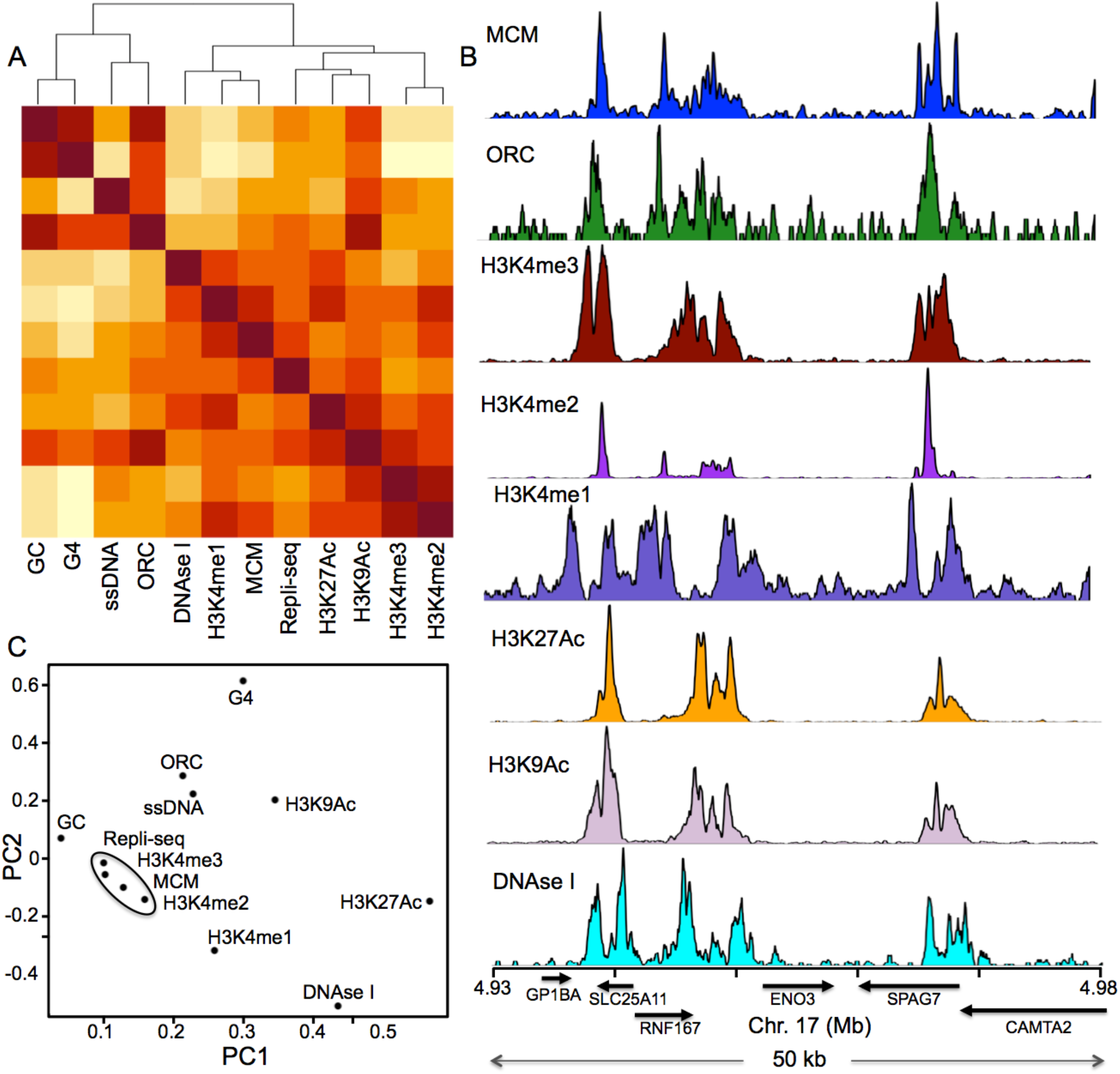
MCM binding in humans is correlated with GC content. (A) Heat map and dendrogram of MCM binding, GC content, and other metrics, as labeled, obtained by hierarchical clustering. (B) Measurements in (A) and (B) displayed for a 50 kb locus on chr17. Data sources cited in main text. (C) Principal component analysis of measurements used in A. 77% and 9% of the variance were accounted for by principal components 1 and 2, respectively.

### Conclusion

Models that account for differences in replication timing have long assumed differences in the propensity of individual licensed origins to fire, with some origins having an intrinsic tendency to fire before others (Dileep & Gilbert, 2018; Rhind & Gilbert, 2013); other models have emphasized the role of stochastic firing of licensed origins in determining replication timing (Arbona, Goldar, Hyrien, Arneodo, & Audit, 2018; Hawkins et al., 2013; Hyrien, 2016; Kelly & Callegari, 2019; Patel, Arcangioli, Baker, Bensimon, & Rhind, 2006; Rhind, 2006). Our study clearly delineates regions, most notably the inactive X chromosome, that are regulated at a post-licensing step. However, we conclude that outside of these regions, most chromosomal differences in replication timing can be attributed to differences in MCM loading along the chromosome.

## Materials and Methods

### Cell culture and media

All *S. cerevisiae* strains were in S288C background. *S. cerevisiae* experiments were carried out using standard YPD (yeast peptone dextrose) medium [2% (wt/vol) glucose, 1% yeast extract, 2% (wt/vol) peptone]. *S. pombe* experiments were carried out in Yeast Extract Dextrose (YES); logarithmically growing cells were arrested in early S-phase by adding 15 mM hydroxyurea and incubating for 3 hours. Strains are listed in Supplementary Data Table 2.

### Tagging Mcm with MNase

Human and mouse MCM2 genes, tagged with micrococcal nuclease at the C-terminus using the same tag that was employed in yeast were cloned into a lentiviral vector. The plx302-mMCM2-MNase was made as follows: Mouse MCM2 cDNA was amplified from plasmid mEmerald-MCM2-N-22, which we received as a gift from Michael Davidson (Addgene plasmid # 54164; http://n2t.net/addgene:54164; RRID:Addgene_54164). 3X FLAG-MNase was amplified from pGZ108 (pGZ108 (pFA6a-3FLAG-MNase-kanMX6), which we received as a gift from Steven Henikoff (Addgene plasmid # 70231; http://n2t.net/addgene:70231; RRID:Addgene_70231). The lentiviral plasmid plx302 was digested with NdeI/NheI and 530bp of plx302 starting from the NdeI cut site was also PCR amplified. All the fragments were assembled by Gibson Assembly as per the manufacturer’s instructions (NEB # E2611S). Finally, to restore the Mcm2 protein sequence to wild type, the Trp 185 residue was changed to Arg using Q5 Quick Mutagenesis Kit (NEB #E0552) according to the manufacturer’s instructions

The plx302-hMCM2-Mnase was made in two stages: First, a Gateway destination vector containing FL-MNase immediately downstream of the attR2 recombination site was constructed. For this, the Gateway destination vector plx302 was linearized with NdeI/NheI, 3XFLAG-MNase was amplified from plx302-mMCM2-MNase and cmR-ccdB was amplified from plx302. These fragments were assembled using Gibson Assembly as before to create plx302-FL-MNase Gateway destination vector. Next, human MCM2 cDNA from plx304-hMCM2-V5 (a gift from Dr. Patrick Paddison, FHCRC) was shuttled to the Gateway donor vector pDONR221 (a gift from Dr. Valeri Vasioukhin, FHCRC) using BP clonase (ThermoFisher Scientific #11789100) as per the manufacturer’s instructions. The resulting entry clone was then used to shuttle hMCM2 cDNA to the plx302-FL-MNase by LR clonase (ThermoFisher Scientific # 1253810) to obtain the final expression vector.

### Primers used for mutagenesis of mouse MCM2

1. XXI-57: GAGCATGGCAGGGCCCAGGCTGGAGATCC
2. XXI-58: ACCCACTCGCGCACCGAGTGGCCCTTGAGG

### Primers used to amplify DNA segments for Gibson cloning of mouse MCM2

1. XXI-51: CTTGGCAGTACATCAAGTGTATCATATGCCAAGTACGCCCCCTATTG
2. XXI-52: GAGAGACTCAGAAGACTCCGCCATCATAGTGACTGGATATGTTGTGTTTTAC These two were used to PCR out CMV promoter
3. XXI-53: GTAAAACACAACATATCCAGTCACTATGATGGCGGAGTCTTCTGAGTCTCTC
4. XXI-54: TTAACCCGGGGATCCGTCGACCGAACTGCTGTAGGATCAGTTTGC These two were used to amplify mouse MCM2 cDNA
5. XXI-55: GCAAACTGATCCTACAGCAGTTCGGTCGACGGATCCCCGGG
6. XXI-56: CTTAACGCGCCACCGGTTAGCGCTACTGGCCGCTATCGGCGTTATC These two were used to amplify FL-MNase

### List of sequencing primers used to confirm sequences of mouse MCM2

1. XXI-59: CATTGACGTCAATAATGACGTATGTTCC
2. XXI-60: TGGCTAACTGTCGGGATCAACAAG
3. XXI-61: TGATGAAGATGTGGAGGAGCTGAC
4. XXI-62: GTTGCTGCAGATCTTTGACGAGGC
5. XXI-63: AGCTGACCGGCATTTACCATAATAAC
6. XXI-64: GTGTCTCATTGACGAGTTTGACAAG
7. XXI-65: CTCAACCAGATGGACCAGGATAAAG
8. XXI-66: CCTGAGAAGGATCTGATGGACAAG
9. XXI-67: GTTCGATAAAGGCCAACGCAC
10. XXI-68: GTTGCGTCAGCAAACACAGTG

### Primers used to amplify DNA segments for Gibson cloning of plx302-FL-MNase Gateway destination vector

1. Plx302_FLMNase_frag1_fwd: TGCCCACTTGGCAGTACATCAAGTGTATCATATGCCAAGTACGCCCCCTATTG
2. Plx302_FLMNase_Frag1 rev: ATCTTTAATTAACCCGGGGATCCGTCGACCAACCACTTTGTACAAG These two were used to PCR CMV-ccdb fragment from plx302.
3. Plx302_FLMNase Frag2 fwd: CGTTTCTCGTTCAGCTTTCTTGTACAAAGTGGTTGGTCGACGGATCCCCGGGTTAATTAAAGATTACAAG
4. Plx302MNase_Frag2 rev: TTGTCGACTTAACGCGCCACCGGTTAGCGCTAGCTCATTACTACTGGCCGCTATCGGCG These two were used to PCR out FL-MNase from plx302-mMCM2-FLMNase

### Sequencing primers used for humanMCM2 plasmid

1. XXI-60: TGGCTAACTGTCGGGATCAACAAG
2. XXI-67: GTTCGATAAAGGCCAACGCAC
3. XXI-68: GTTGCGTCAGCAAACACAGTG

### Human and mouse MCM2-FLAG-MNase cell lines

Cells were transduced with lentiviral supernatant containing plx302-MCM2-FL-MNase, infected cells were selected out on Puromycin (2.5μg/ml) and expanded for further experiments. The expression of tagged MCM2 in HeLa and Patski cells was confirmed by Western blot using anti-MCM2 rabbit polyclonal antibody (HPA031495 Sigma) at 1:1,000 dilution.

### Yeast ChEC protocol

In budding yeast, ChEC-seq was carried out as previously described (Foss et al., 2019). Fission yeast were transferred from log phase to 15 mM HU for 3 hours, and then processed as in budding yeast.

### Mammalian ChEC protocol

Cells were grown in 6 well plates until >100% confluent, trypsinized and washed with a Wash Buffer (20mM HEPES, 110mM Potassium Acetate, 5mM Sodium Acetate, pH to 7.3 with NaOH). Cells were permeabilized with 0.02% digitonin in the Wash Buffer along with protease inhibitors (Roche # 04693159001) for 5 mins at RT. CaCl_2_ was added to a final concentration of 2.5mM for differing amounts of time, as indicated. The MNase activity was stopped by adding equal volume of a 2X Stop Buffer (400mM NaCl, 20mM EDTA, 4mM EGTA, 2% SDS). Proteinase K was added to a final concentration of 0.4mg/ml and samples incubated for 1 hr at 50° C. DNA was isolated using standard Phenol:Chloroform extraction and air dried DNA pellets resuspended in 0.1X TE (pH 8). All samples were treated with RNase A (0.3 μg/ml) before being processed for library preparation.

### Sequencing

Sequencing was performed using an Illumina HiSeq 2500 in Rapid mode employing a paired-end, 50 base read length (PE50) sequencing strategy. Image analysis and base calling was performed using Illumina’s Real Time Analysis v1.18 software, followed by ‘demultiplexing’ of indexed reads and generation of FASTQ files, using Illumina’s bcl2fastq Conversion Software v1.8.4.

### Sequence analysis

fastq files were aligned to genome assemblies for Saccharomyces cerevisiae (sacCer3), Schizosaccharomyces pombe (ASM294v2), homo sapiens (GRCh38/hg38), or mus musculus (mm10) using bwa alignment software (H. Li & Durbin, 2009) with the −n 1 option, which causes reads that map to more than one location to be randomly assigned to one of those locations. bam files were then processed with Picard’s CleanSam, SortSam, FixMateInformation, AddOrReplaceReadGroups and ValidateSamFile tools (http://broadinstitute.github.io/picard/). Per-base pair read depths were then determined with BedTools’ genomecov tool (Quinlan & Hall, 2010).

### Analysis of the hybrid *M.musculus/M.spretus* Patski cell line

A hybrid musculus-spretus genome was generated in silico by (1) identifying all sites in which the musculus (mm10) and spretus (SPRET_EiJ) genomes differed by a single nucleotide substitution and (2) replacing all such sites in the mm10 genome with the corresponding SPRET_EiJ sequence. fastq files were then aligned to the mm10 genome and the hybrid mm10 - SPRET_EiJ genome using the bwa alignment software (H. Li & Durbin, 2009) and processed with Picard’s tools as described above. Reads were retained only if they (1) contained at least one of the single-nucleotide differences that were used to generate the hybrid mm10 - SPRET_EiJ genome and (2) had no mismatches with the genome to which they were aligned. Reads that did not match these criteria were filtered out using Picard’s FilterSamReads tool (http://broadinstitute.github.io/picard/). Per-base pair read depths were then determined with BedTools’ genomecov tool (Quinlan & Hall, 2010), with those coming from the filtered bam files generated from the mm10 and hybrid genome alignments constituting the read depths assigned to the *musculus* and *spretus* genomes, respectively.

### Replication Simulations

The replication simulations had 3 parameters: (1) the bin size into which MCM2-ChEC will be grouped to generate probability distributions; (2) the number of MCM complexes that are activated at the beginning of S phase; and (3) the distance replication forks are allowed to go before another batch of MCM complexes, which will correspond to the number of converged forks, will be activated. For these parameters, we chose 1 kb, 300, and 5 kb, respectively. The second parameter can be thought of as reflecting the number of MCM complexes in the cell, and the third parameter can be thought of as reflecting the frequency with which replication factors are “recycled” after diffusion. Replication simulation results were very robust to variation in these parameters, including completely eliminating “recycling”.

Per-base pair read depths from *cerevisiae* MCM2-ChEC data sets were binned in 1 kb bins across the genome and these numbers were used to generate probability distributions. 300 MCM complexes were chosen from this probability distribution, the central base pair from each 1 kb bin was assigned the number 0, base pairs to the right of this base pair were assigned the numbers +1, +2, +3, etc., and base pairs to the left were assigned corresponding negative numbers. These numbers increase in absolute value until they either (1) run off the end of the chromosome or (2) converge with numbers coming from another activated MCM complex. At this point, every base pair in the genome will have been assigned a number, which corresponds to the order in which it has been “replicated”, and we then count the number of replication forks that would have converged and activate that number of MCM complexes, chosen at random from the original probability distribution after excluding (1) MCM complexes that have already been activated and (2) MCM complexes that have been replicated. This process is repeated until the entire genome has been replicated by a replication fork that has not traveled more than 5 kb. 1000 “cells” were simulated and from these, the replication pattern of the corresponding population was inferred.

### Heat Map and PCA

Heat maps showing correlation coefficients between MCM2-ChEC signals and various parameters, shown in figure 4A, were generated with the “heatmap” function provided in base R, using the default parameters for distance. Principal Component Analysis was done using the “prcomp” function in base R.

### GM12878 repli-seq replication, DNase-hypersensitivity, and histone modifications

We downloaded fastq files from the ENCODE portal (Davis et al., 2018) (https://www.encodeproject.org/) with the following identifiers: ENCFF001GNU, ENCFF001GNY, ENCFF001GOB, ENCFF001GOG, ENCFF001GOM, ENCFF001GOO, ENCFF000BBP, ENCFF000BBQ, ENCFF000BBT, ENCFF000BBW, ENCFF000BBY, ENCFF000BBZ, ENCFF000BCE, ENCFF000BCI, ENCFF000BCT, ENCFF000BCU, ENCFF000BCZ, ENCFF000BDA, ENCFF000BDH, ENCFF000BDN, ENCFF001DDE, ENCFF001DDF, ENCFF001FKI, ENCFF001FKM, ENCFF001FLD, ENCFF001FLG, ENCFF002ERG, ENCFF002ERH, ENCFF002ERJ, ENCFF002ERL, ENCFF248WCN, ENCFF377VCZ, ENCFF485LYA, ENCFF588FTN, ENCFF625VRT. These were then aligned and processed as described above.

## Acknowledgements

We are grateful to the ENCODE Consortium and the laboratories of John Stamatoyannopoulos and Bradley Bernstein for generating the repli-seq, DNase-hypersensitivity, and histone modification data sets for GM12878 and HeLa cell lines. This work was supported by the National Institute of Health Grants R01-GM117446 and T32 CA009657.

## Author contributions

EJF, SS, TGS, HK, AT, UL and AB designed the experiments. SS, TGS, HK, AT and UL carried out the experiments under the supervision of AB. EJF and AB analyzed the data and wrote the manuscript.

---

**Supplementary data figure 1.**
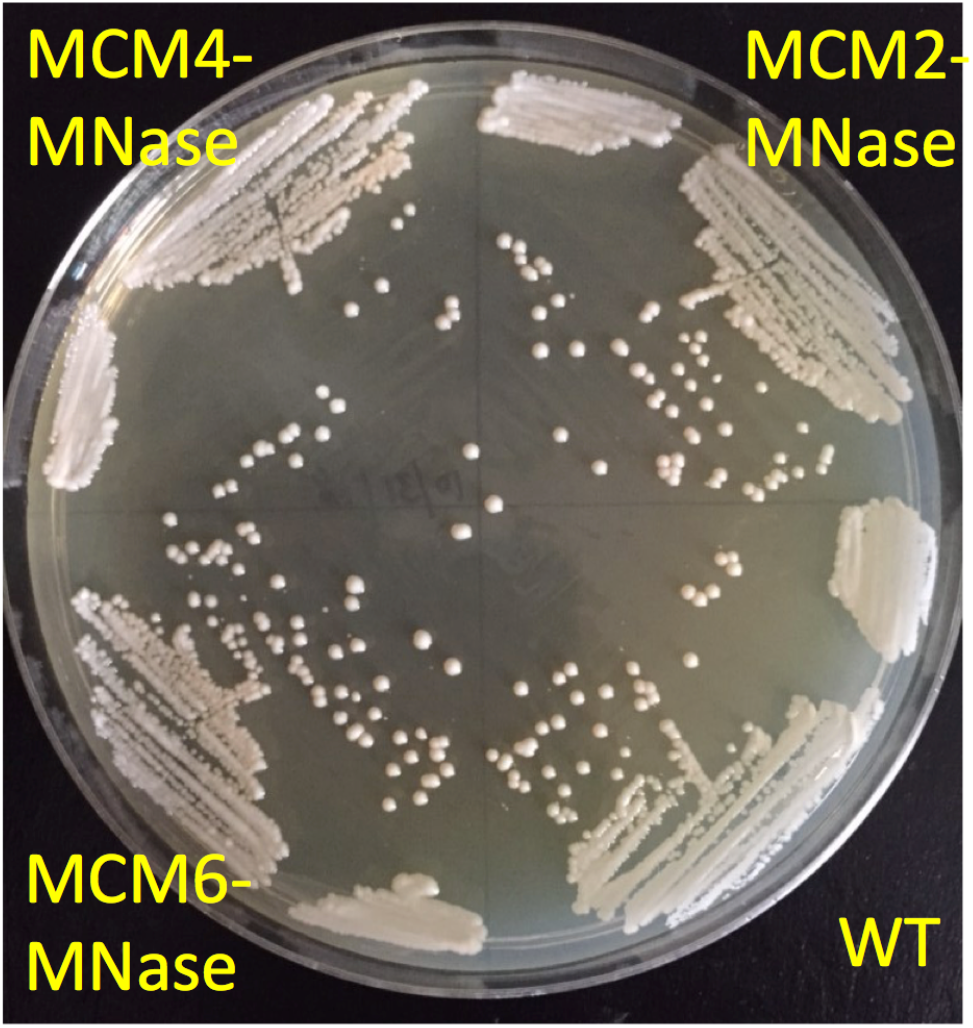
Streaking out strains on rich medium (YEPD) indicates that tagging MCM2, MCM4 and MCM6 with MNase did not affect growth rates.

**Supplementary data figure 2.**
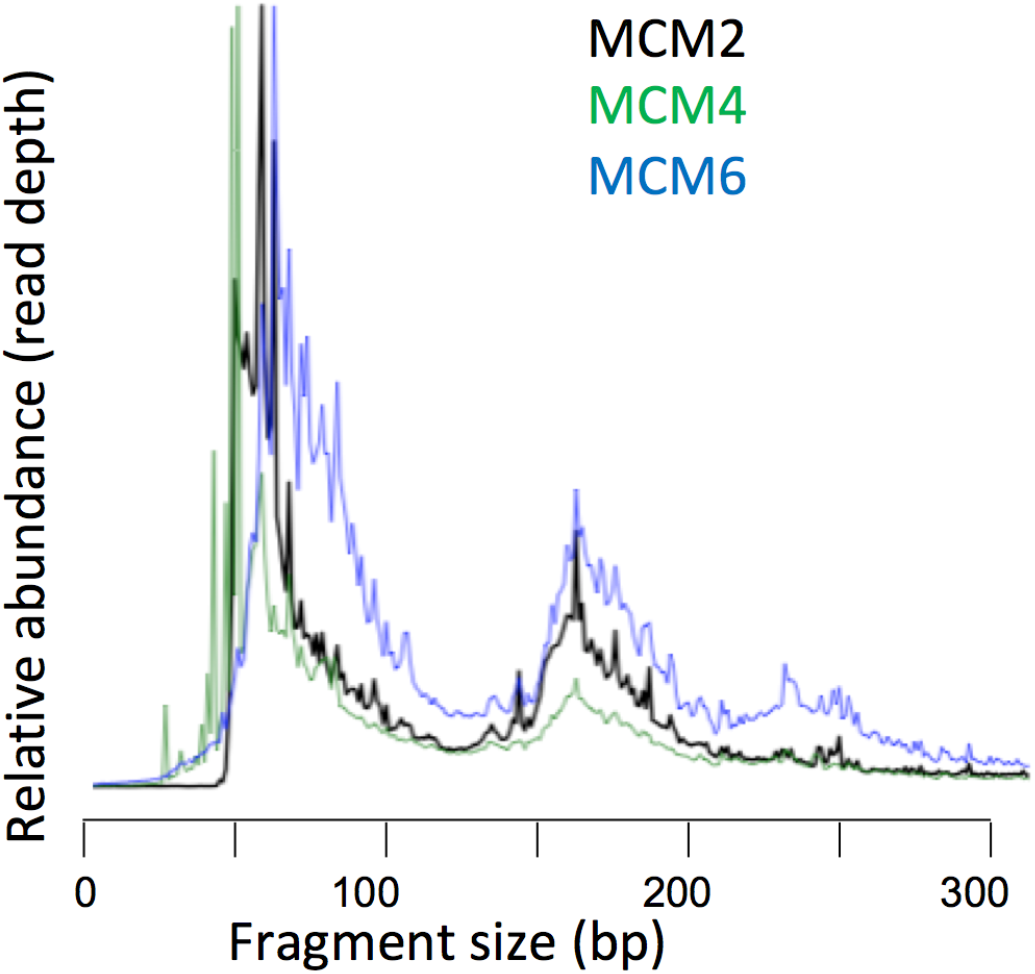
Fragment sizes in libraries from *S. cerevisiae* strains with tagged Mcm2, Mcm4 and Mcm6 are comparable and consistent with the 62 bp of DNA shown to be protected by the MCM helicase complex by cryo-EM.

**Supplementary data figure 3.**
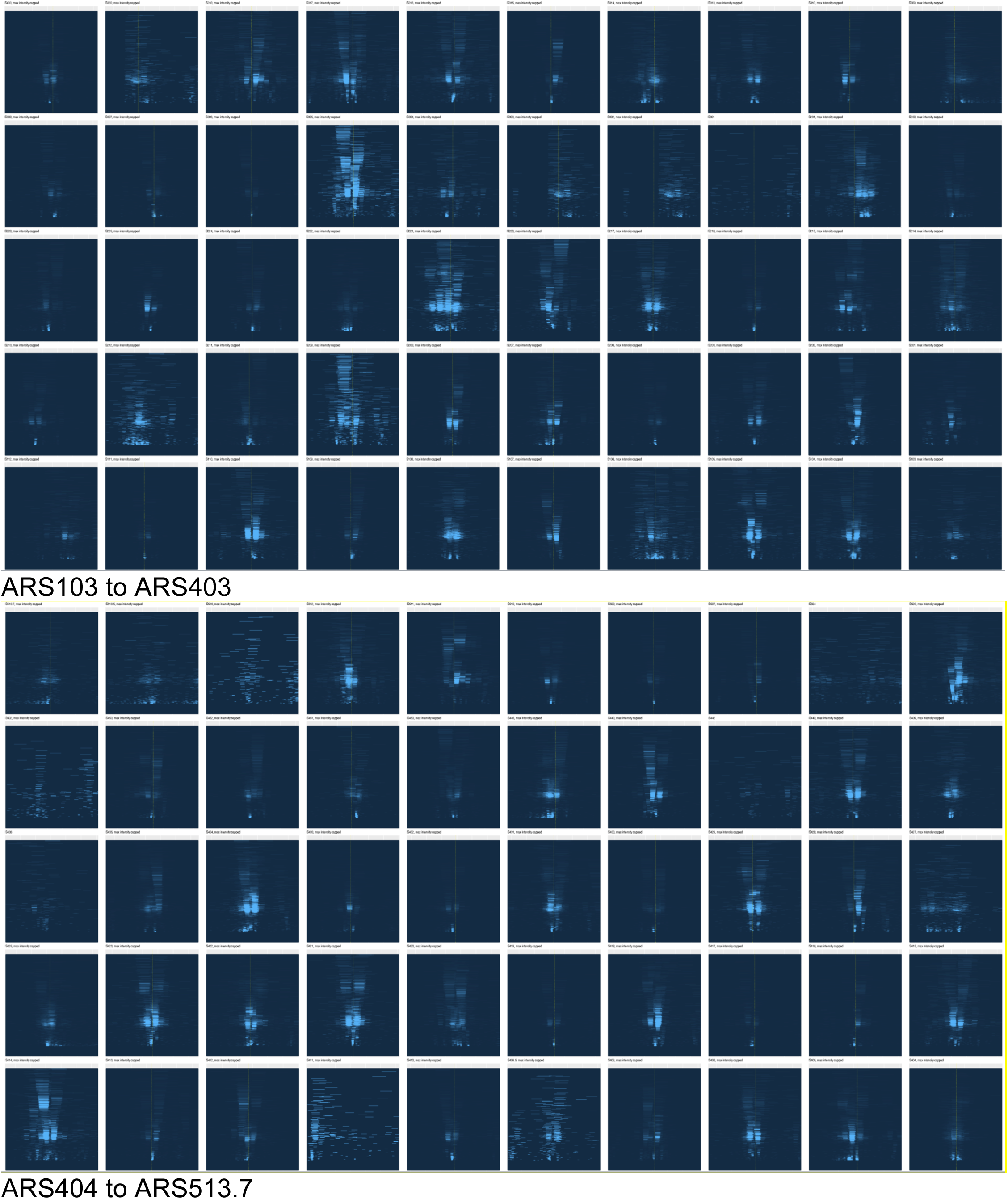
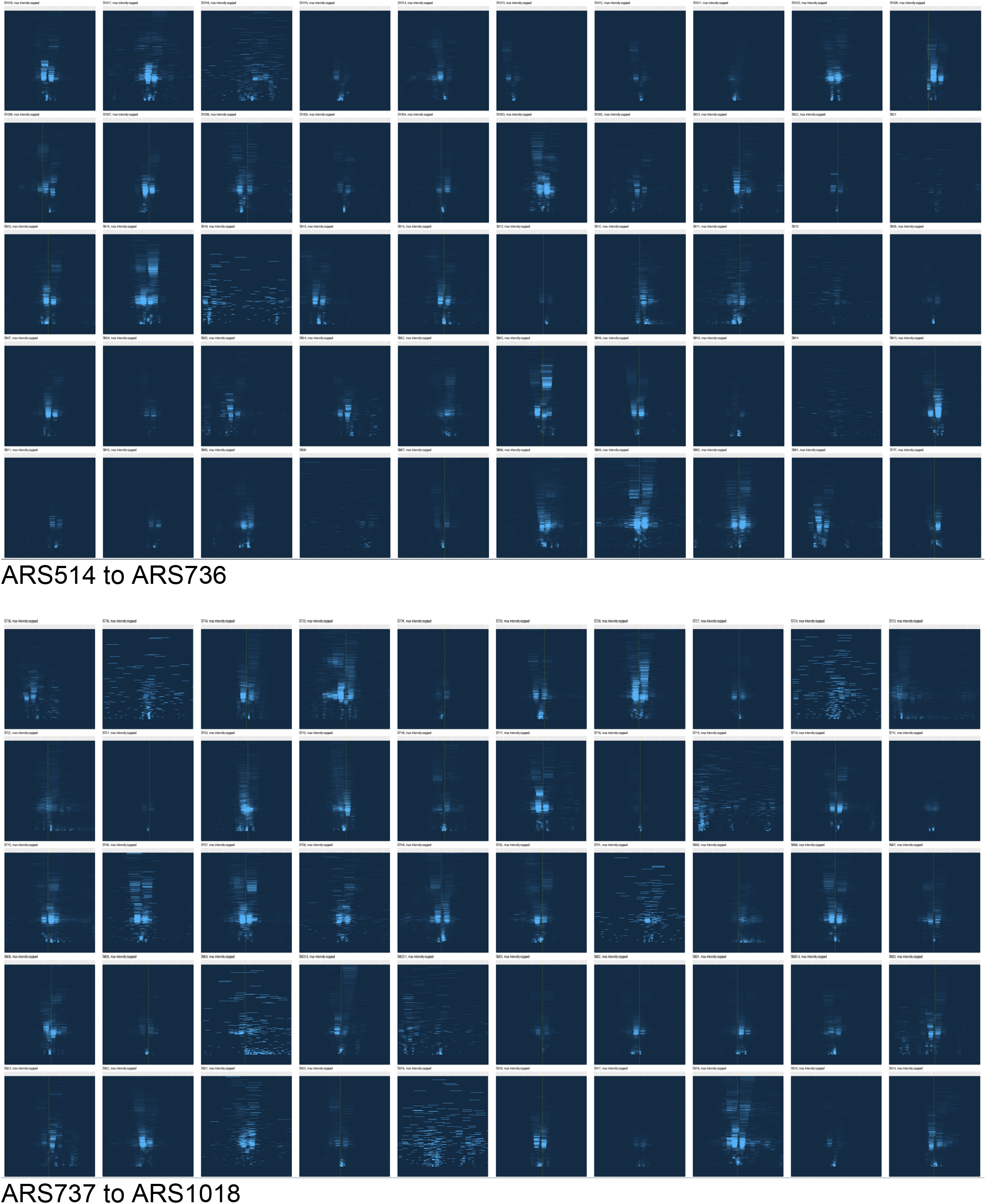
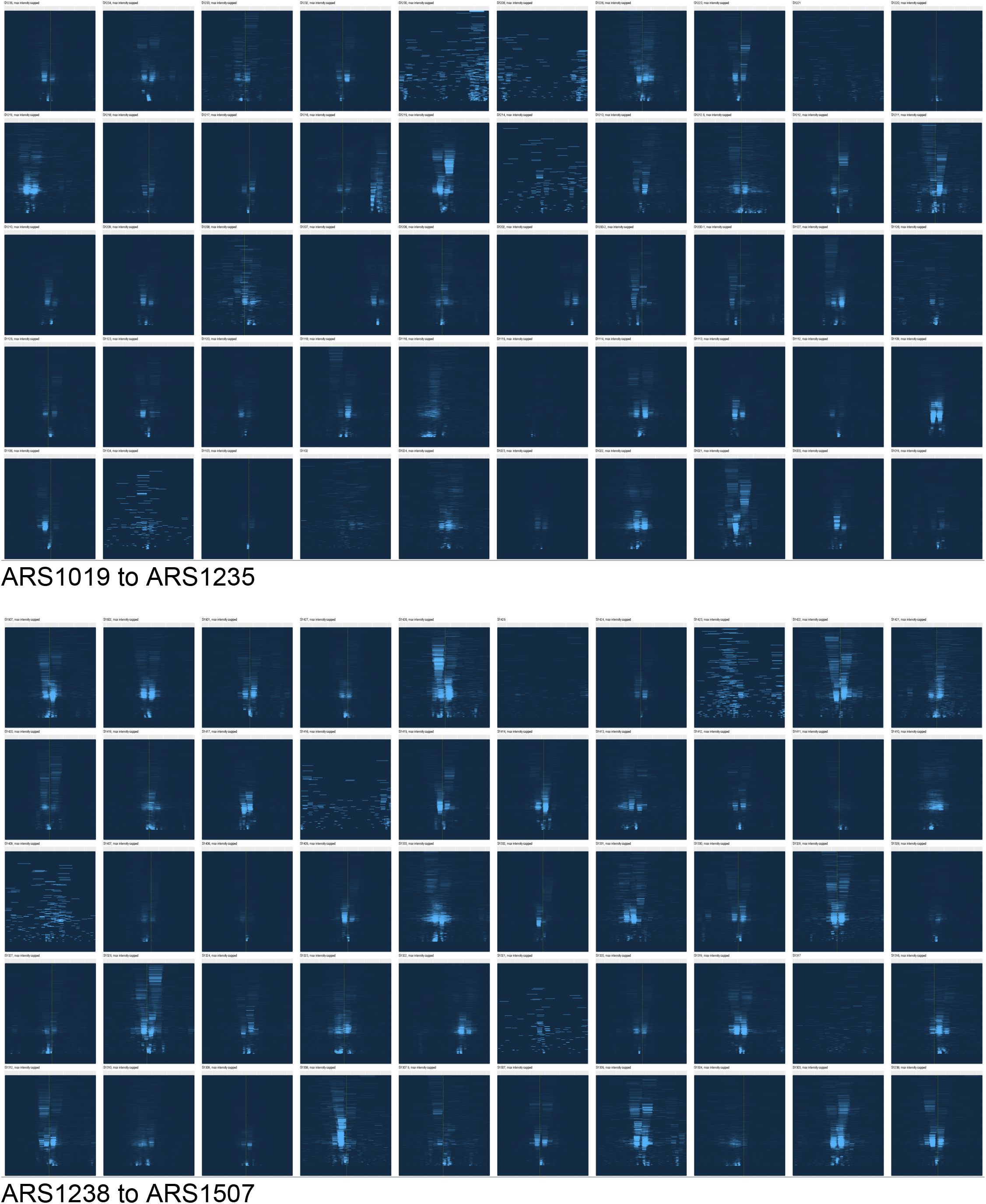
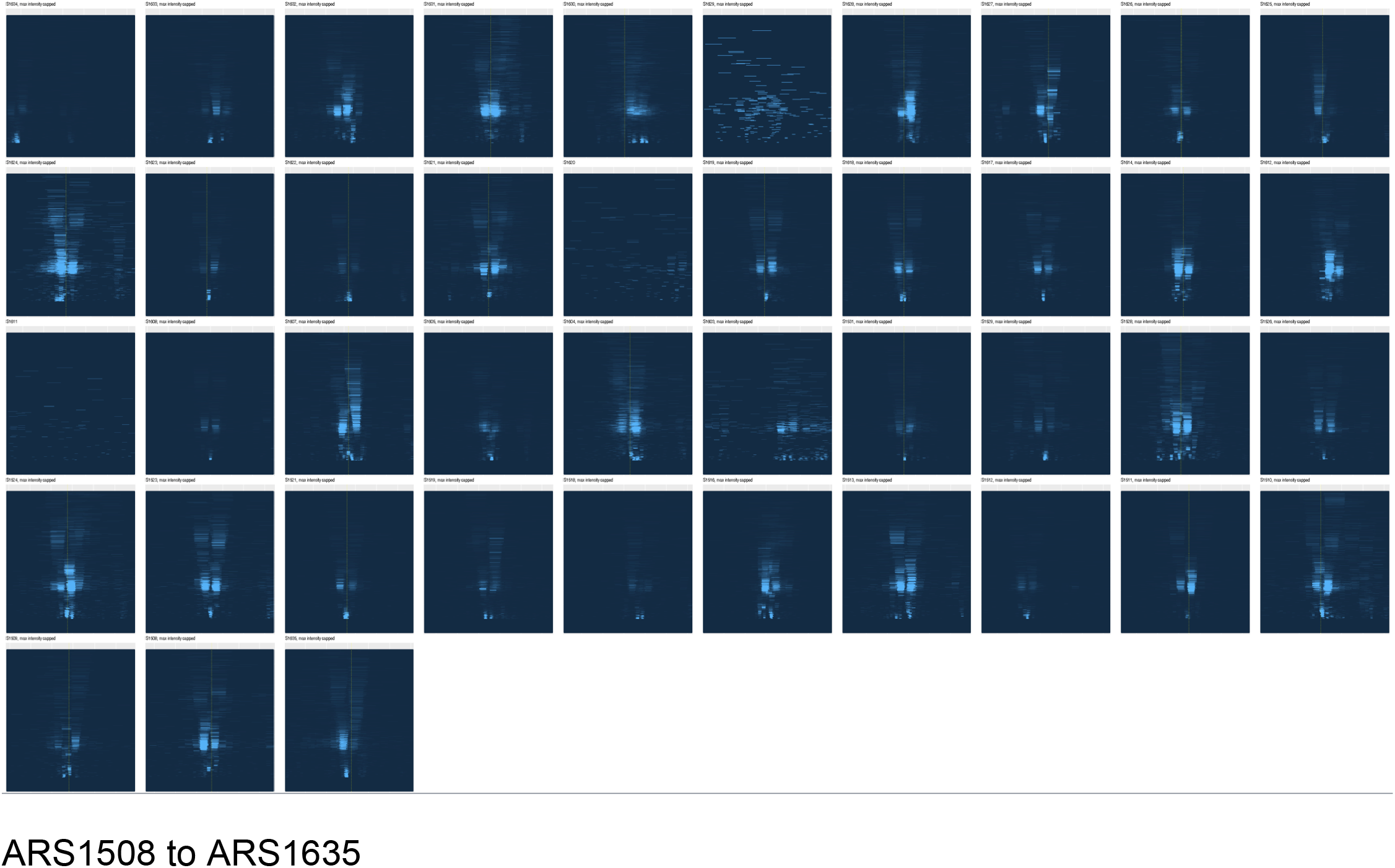
343 origins from Saccharomyces Genome Database (SGD) presented as heat maps. 9 origins listed in SGD were omitted from this analysis due bioinformatic difficulties, such as ambiguous mapping locations. 187 of the remaining 343 origins contained ACS sequences, and these are indicated with dotted lines. Each figure shows a 3 kb span on the X axis. Fragment sizes range from 0 to 500 bp on the Y axis.

**Supplementary data figure 4.**
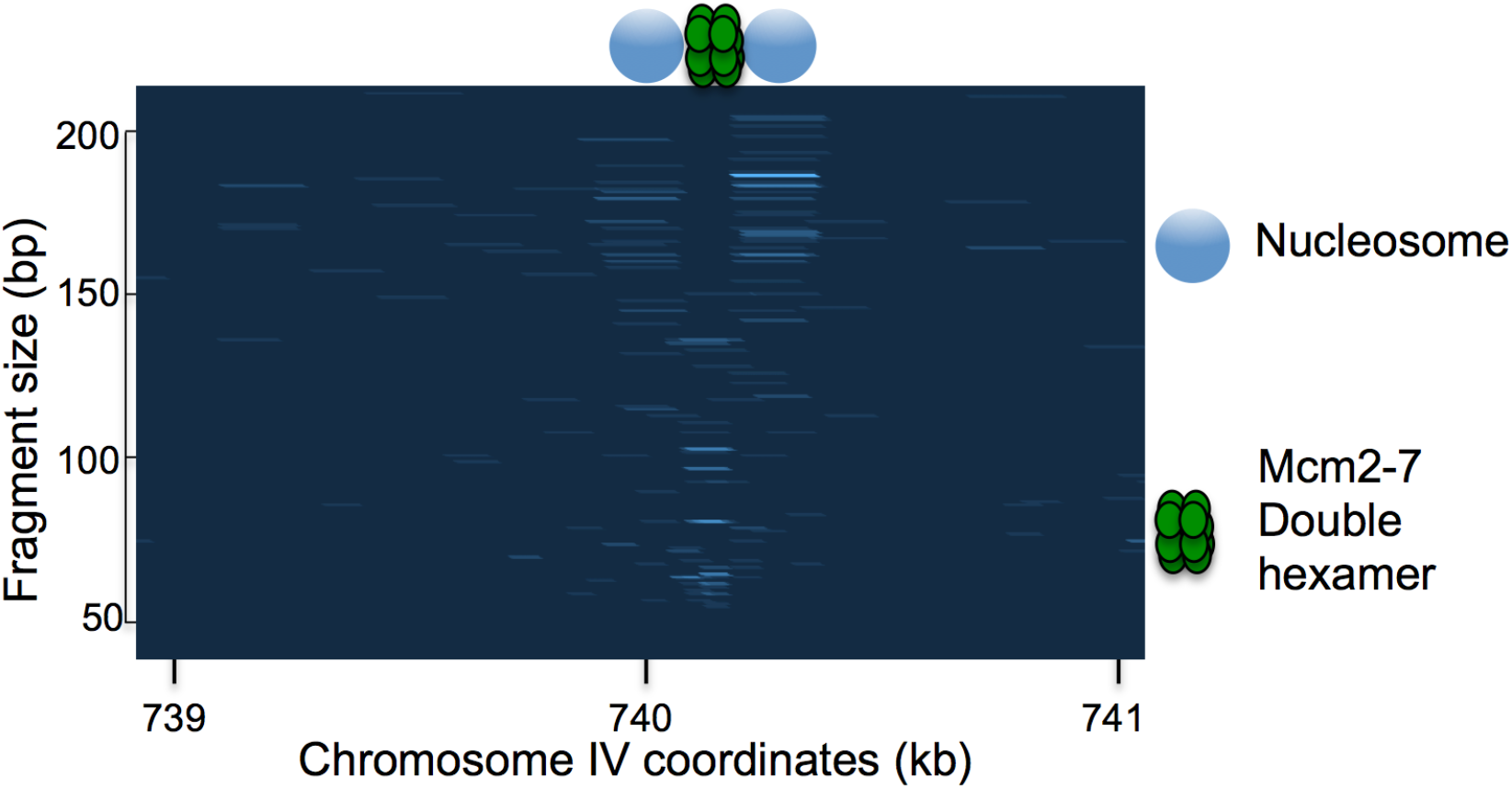
Typical low abundance MCM footprint outside of known origins in *S. cerevisiae*.

**Supplementary data figure 5.**
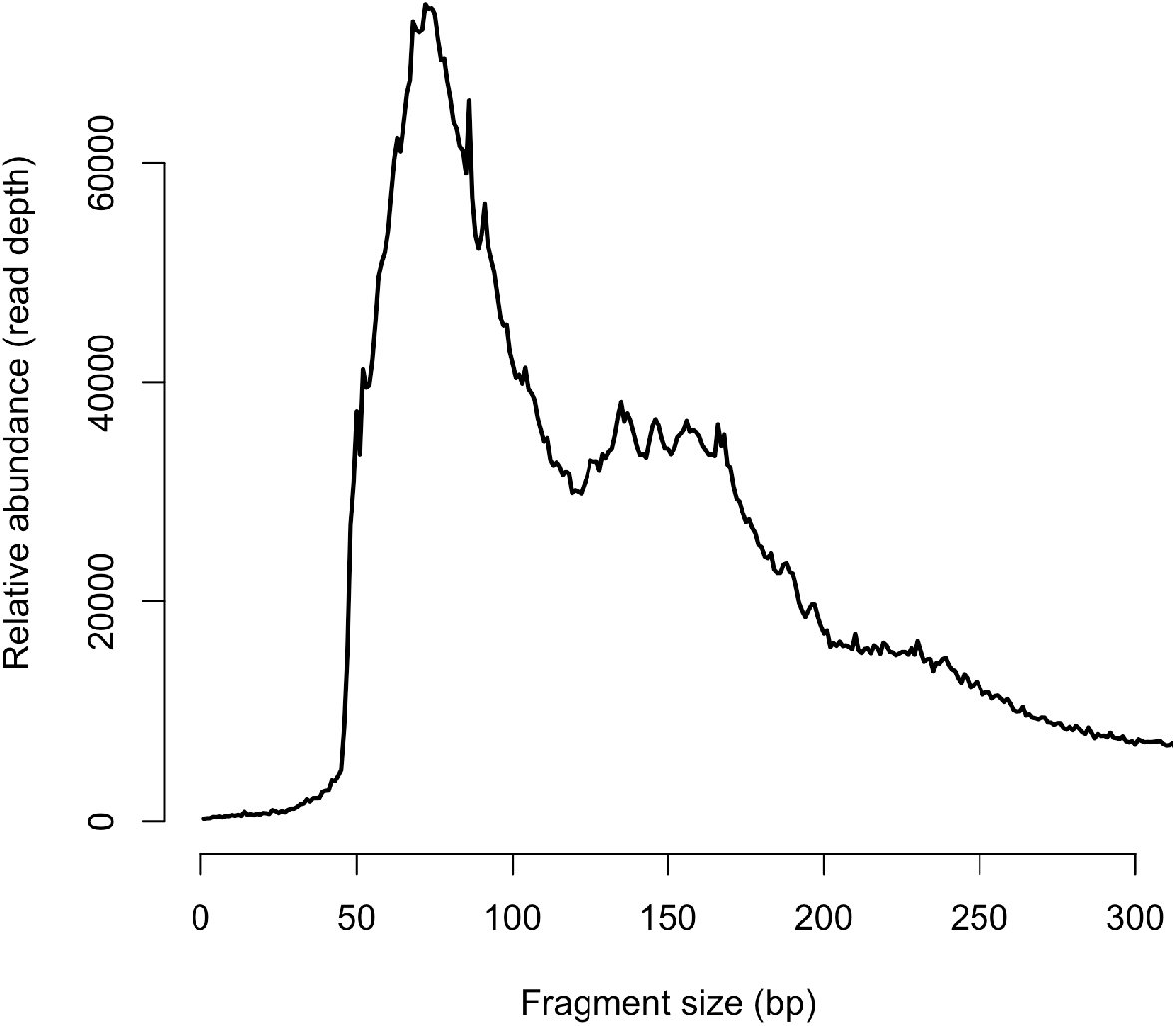
Fragment size distributions for Mcm2-MNase libraries prepared from fission yeast.

**Supplementary data figure 6.**
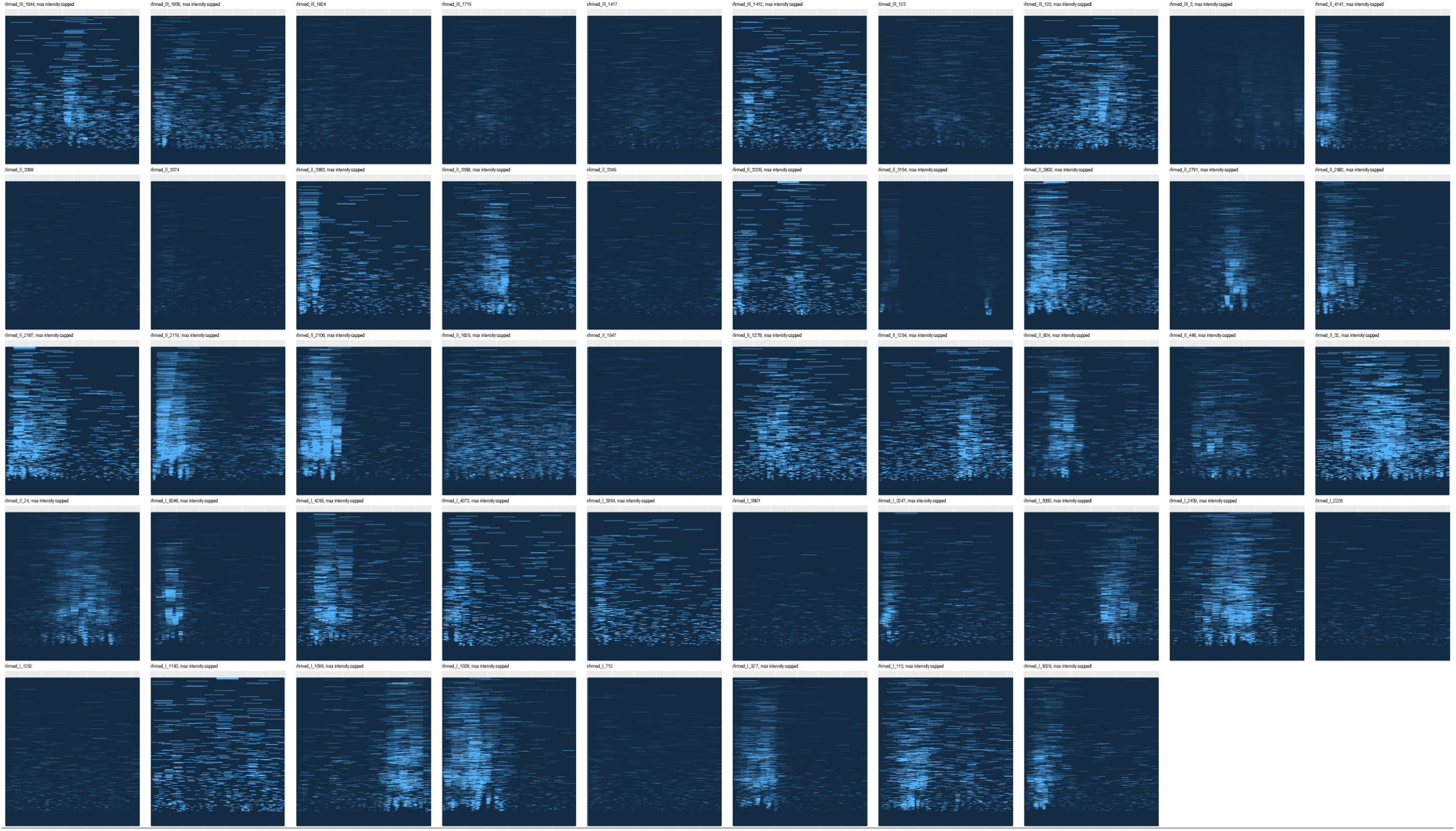
All “confirmed” *S. pombe* origins from OriDB presented as heat maps. Each figure shows a 3 kb span on the X axis. Fragment sizes range from 0 to 500 bp on the Y axis.

**Supplementary data figure 7.**
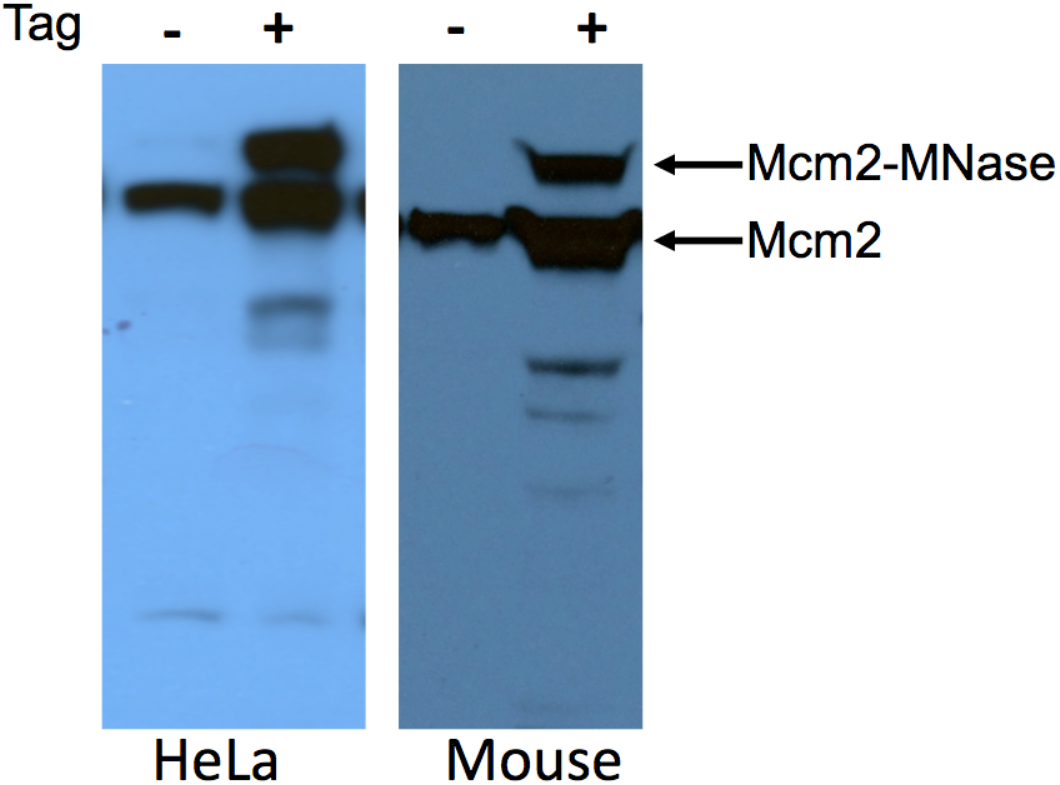
Immunoblot showing expression level of MCM-MNase fusion protein in HeLa and mouse hybrid *M.musculus/M.castaneus* Patski cells.

**Supplementary data figure 8.**
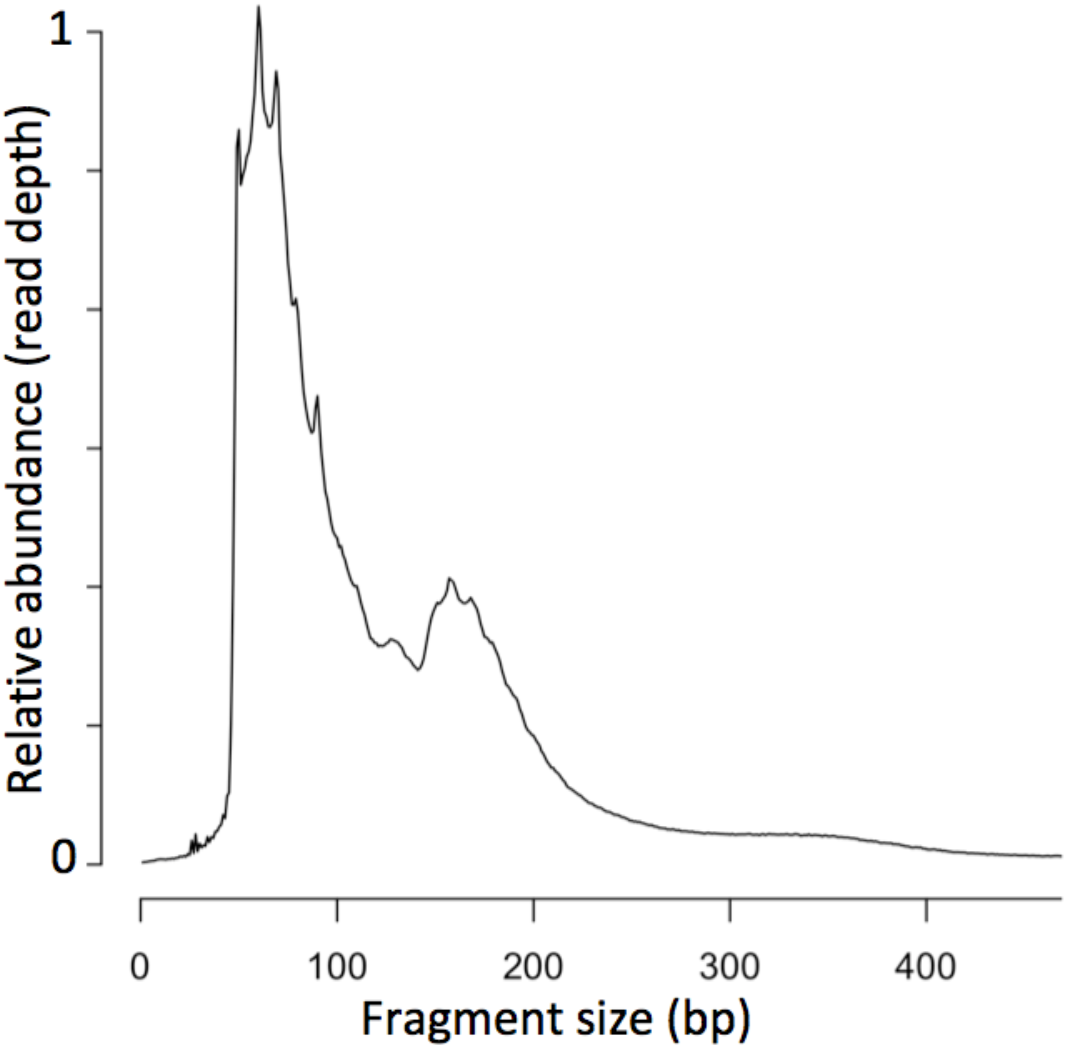
Size distribution of Mcm-ChEC fragments in HeLa cells.

**Supplementary data figure 9.**
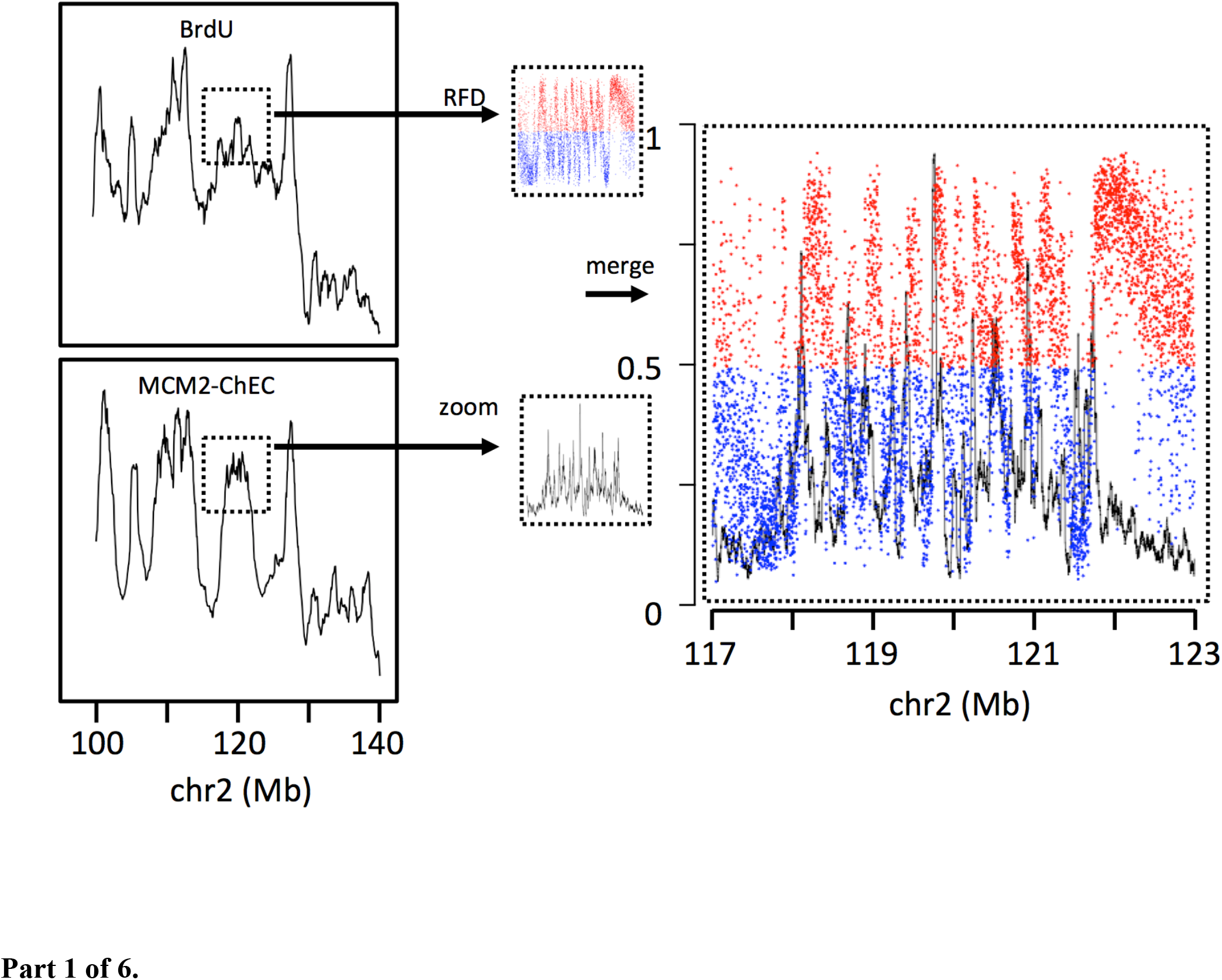
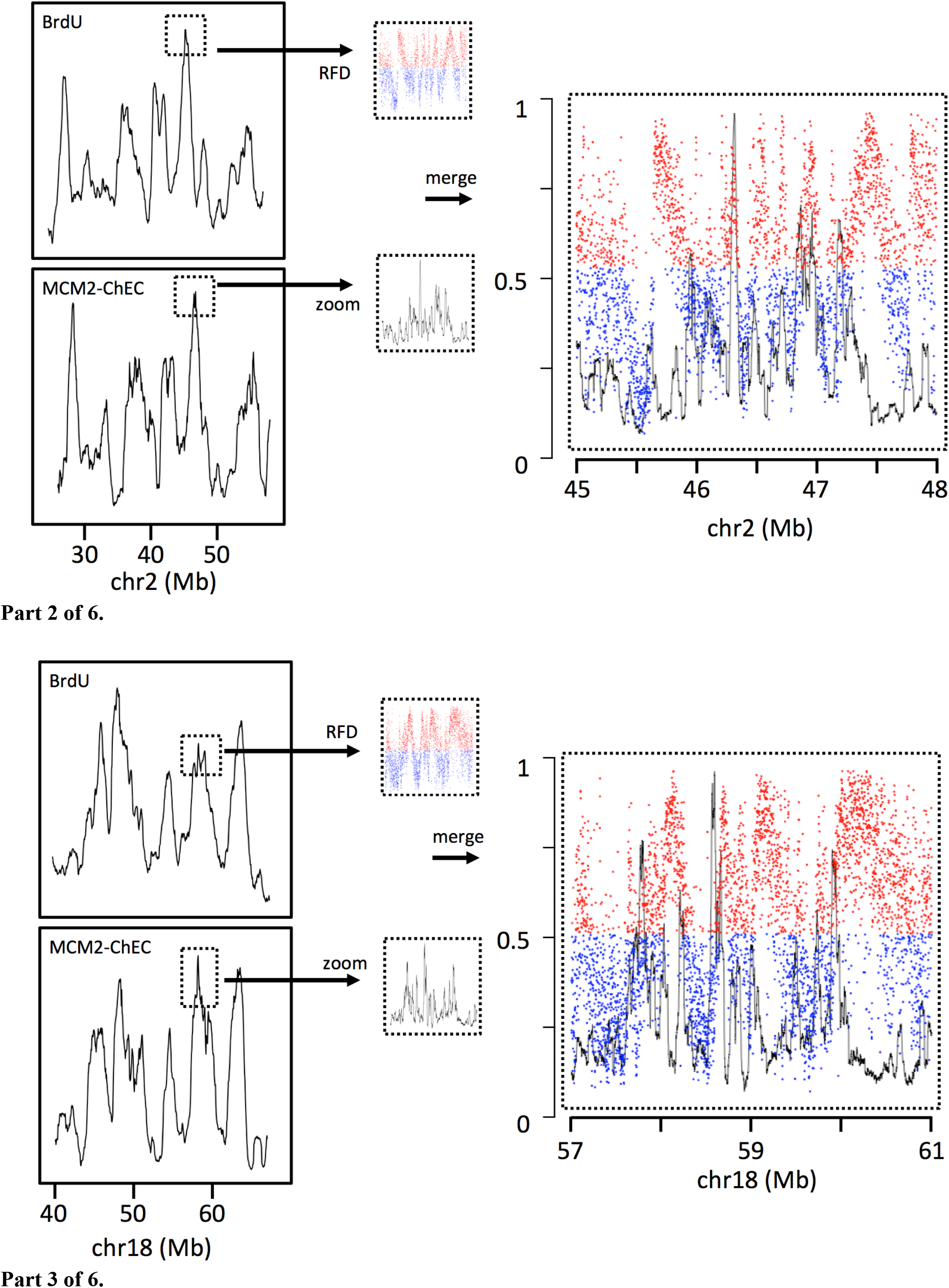
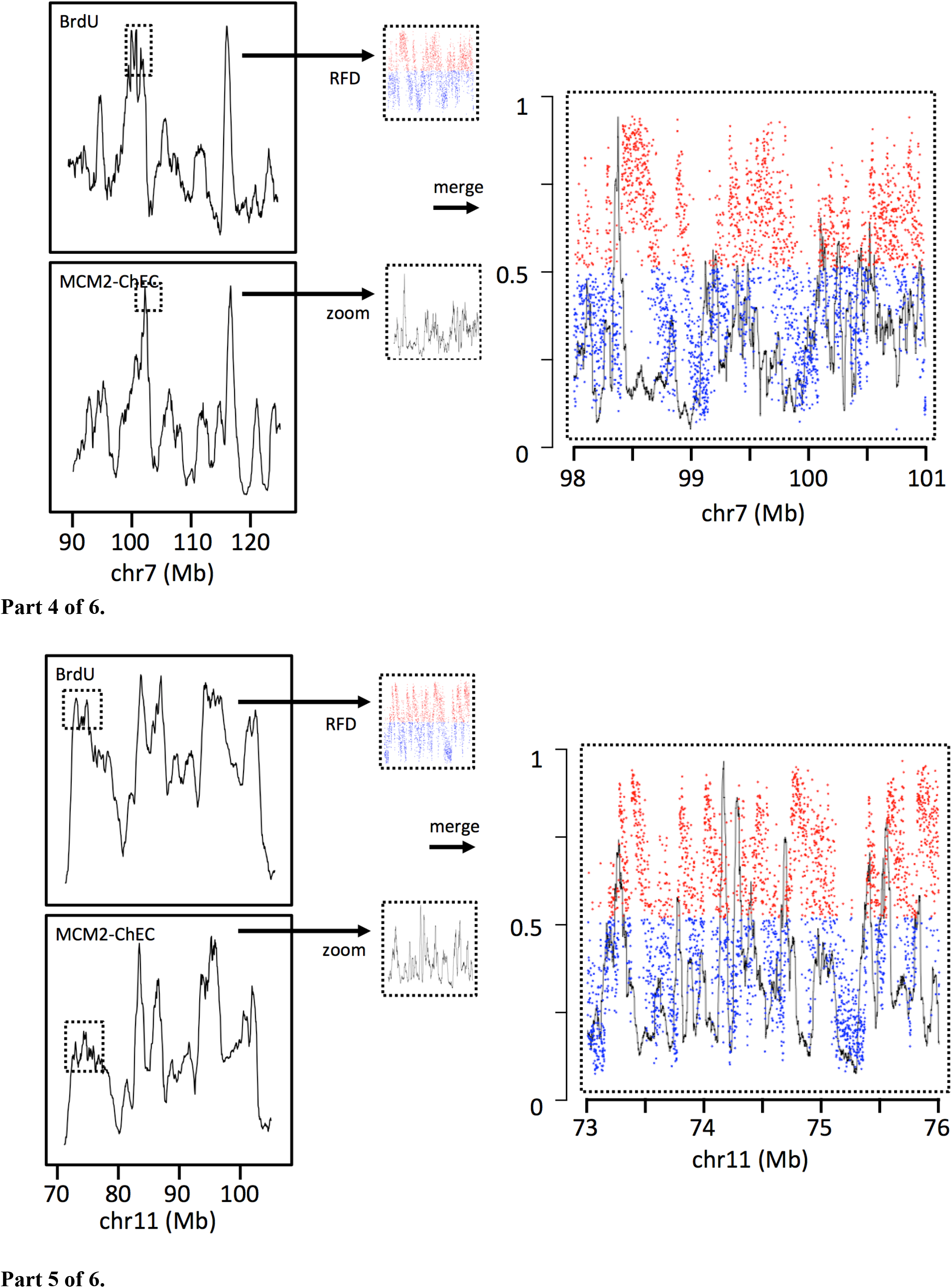
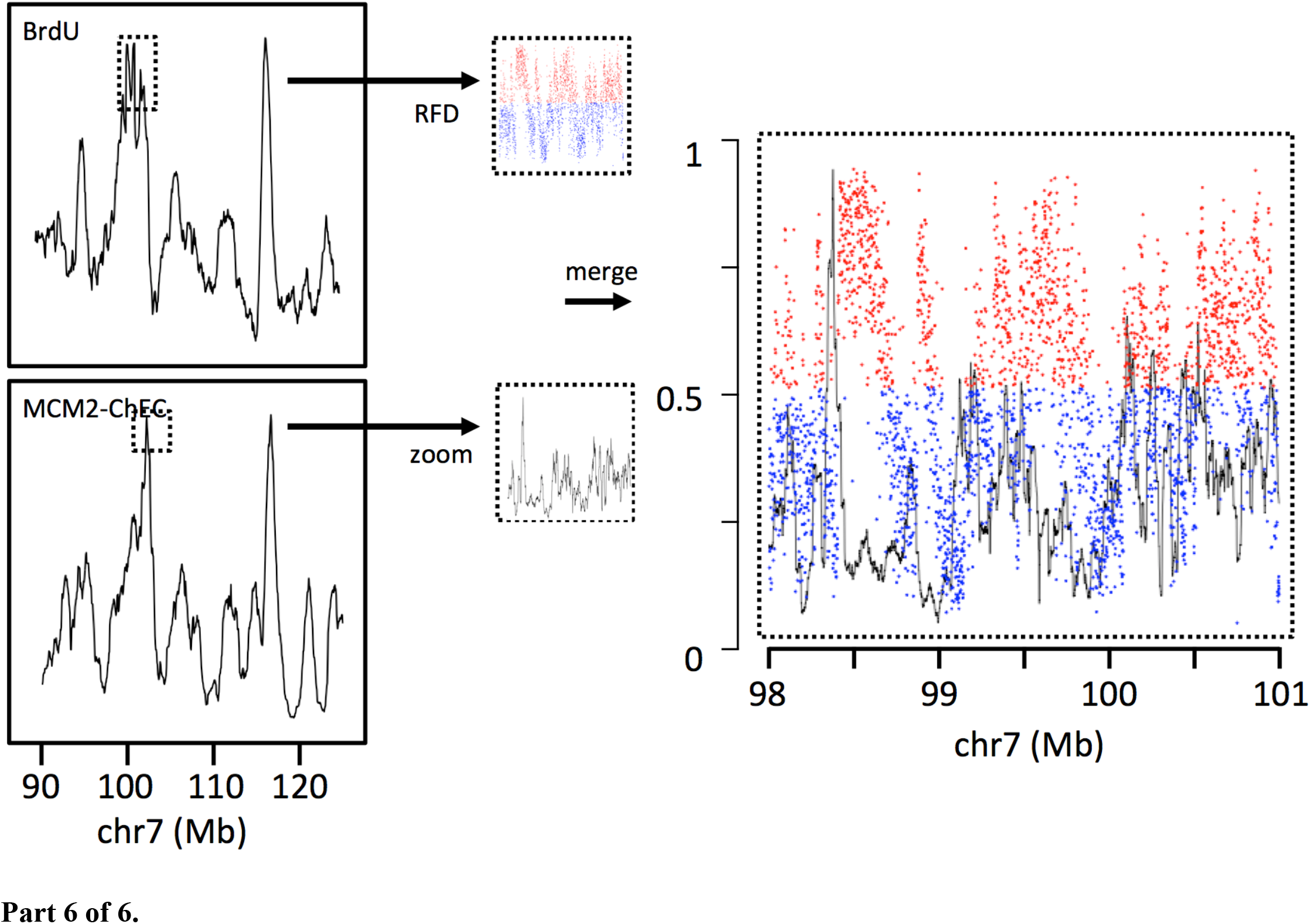
Six examples of replication fork direction assays showing fraction of forks that synthesize the Watson strand that are moving rightward (red and blue dots used to indicate direction of movement of majority of forks) juxtaposed with MCM binding (black) as in Fig. 2B in the main text.

**Supplementary data figure 10.**
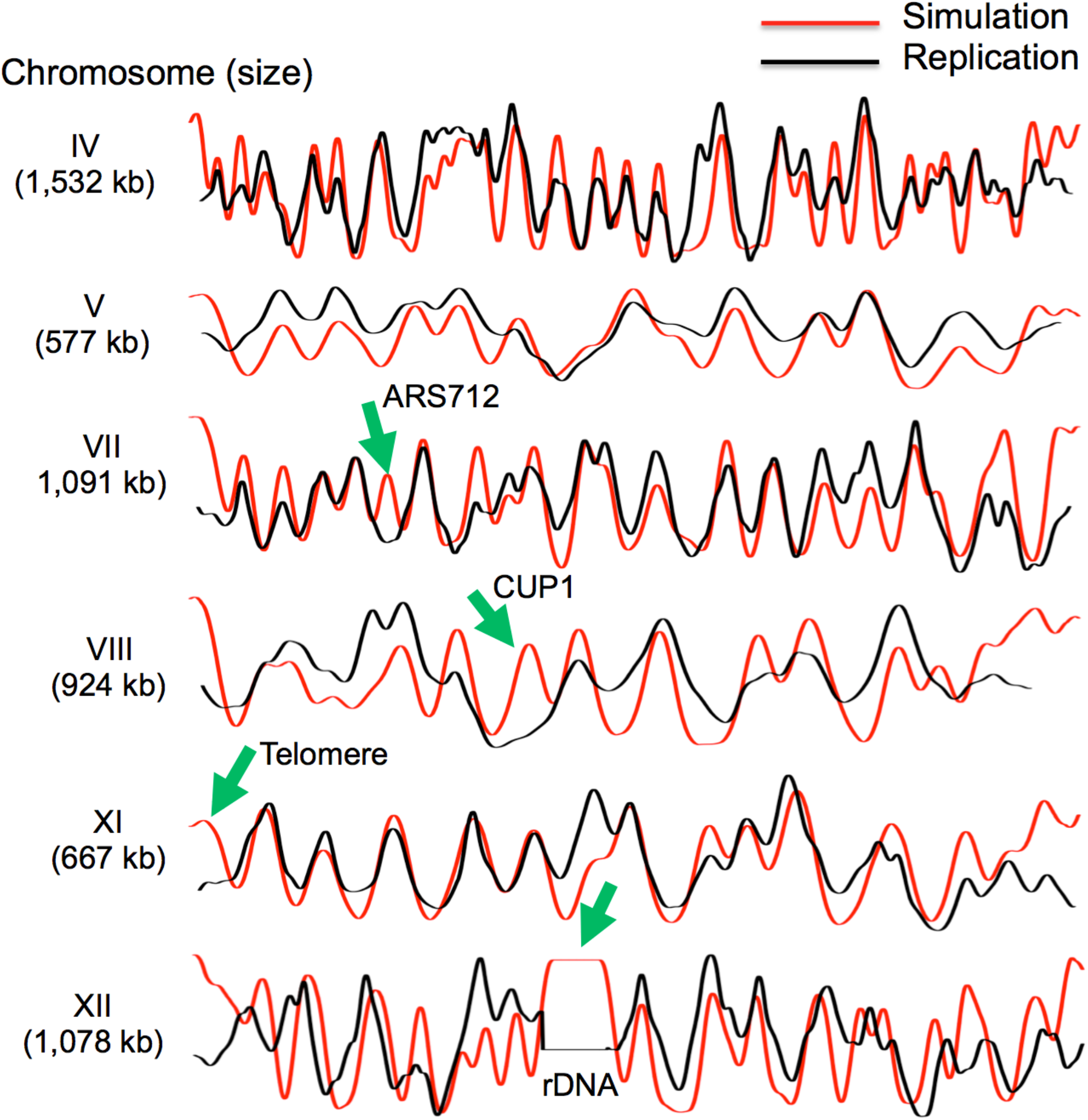
Comparison of replication simulations (red) and replication as assessed by read depth ratios of S phase to G1 phase flow-sorted cells (Yabuki, Terashima, & Kitada, 2002) for seven *S. cerevisiae* chromosomes. Green arrows indicate sites that replicate later than would be predicted based on simulations including repetitive *rDNA* repeats, telomeres *CUP1* locus and *ARS712*. Delayed activation of these well-licenced origins indicates their control at the level of origin firing. *ARS712*, which has been shown to contain both ORC and MCM in ChIP studies (Wyrick et al., 2001), is not active during unperturbed replication, but is active in *rad53* mutants subjected to hydroxyurea (Alvino et al., 2007).

**Supplementary data figure 11.**
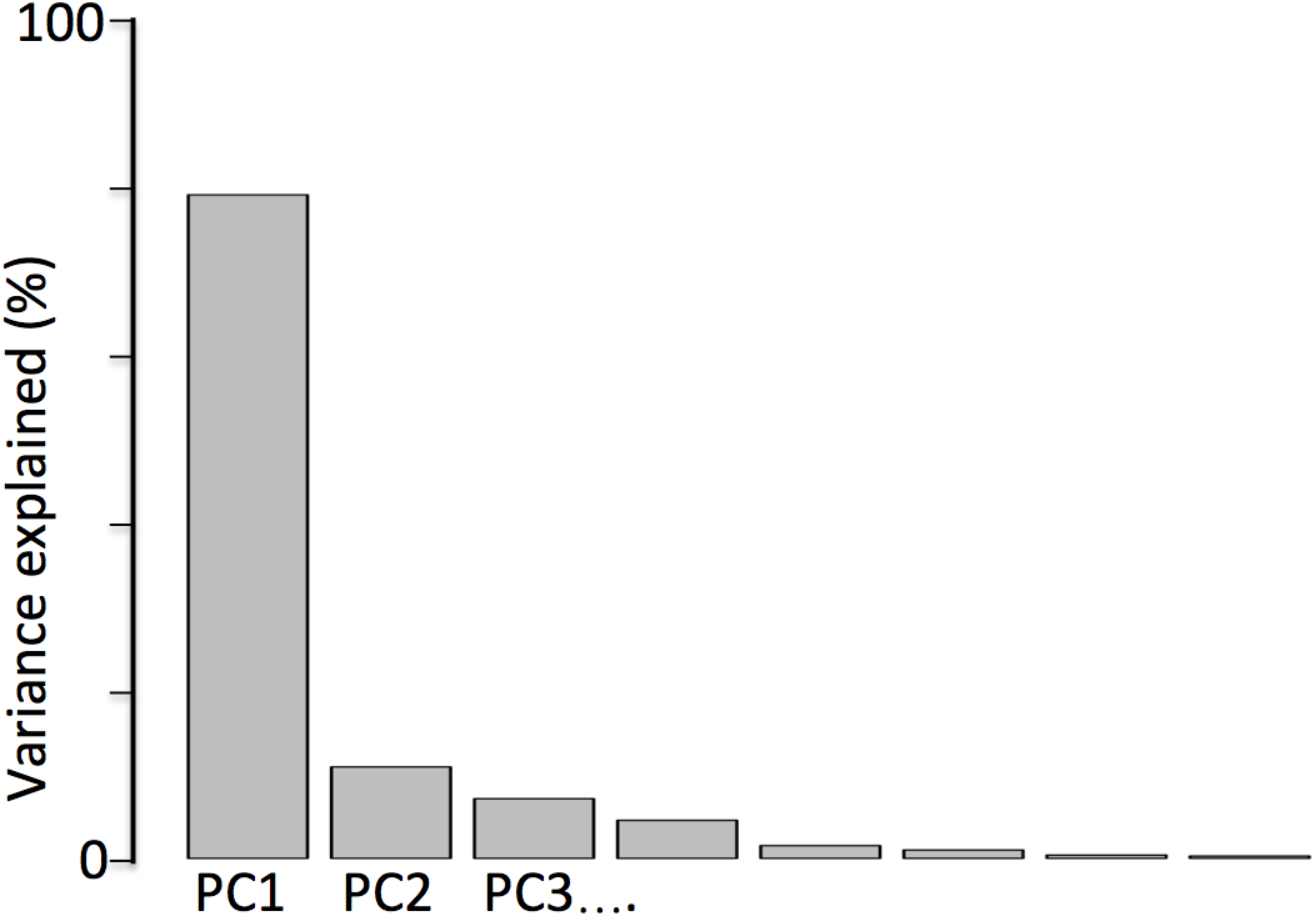
“Scree plot” showing variance accounted for by different principal components in Fig. 4B.

**Supplemenatry data table 1.** Correlation coefficient r values for all pairs shown in heat map in Fig 4A.

|  | MCM | GC | ssDNA | G4 | BrdU | DNase | H3K27Ac | H3K4me1 | H3K4me2 | H3K4me3 | H3K9Ac | Orc |
| --- | --- | --- | --- | --- | --- | --- | --- | --- | --- | --- | --- | --- |
| MCM | 1 | 0.59 | 0.66 | 0.46 | 0.83 | 0.91 | 0.72 | 0.87 | 0.8 | 0.68 | 0.73 | 0.7 |
| GC | 0.59 | 1 | 0.64 | 0.93 | 0.72 | 0.57 | 0.65 | 0.48 | 0.48 | 0.47 | 0.81 | 0.93 |
| ssDNA | 0.66 | 0.64 | 1 | 0.48 | 0.67 | 0.61 | 0.6 | 0.56 | 0.67 | 0.7 | 0.79 | 0.81 |
| G4 | 0.46 | 0.93 | 0.48 | 1 | 0.63 | 0.49 | 0.65 | 0.4 | 0.34 | 0.35 | 0.74 | 0.83 |
| BrdU | 0.83 | 0.72 | 0.67 | 0.63 | 1 | 0.78 | 0.76 | 0.74 | 0.71 | 0.66 | 0.78 | 0.78 |
| DNase | 0.91 | 0.57 | 0.61 | 0.49 | 0.78 | 1 | 0.85 | 0.95 | 0.84 | 0.73 | 0.78 | 0.67 |
| H3K27Ac | 0.72 | 0.65 | 0.6 | 0.65 | 0.76 | 0.85 | 1 | 0.85 | 0.79 | 0.76 | 0.89 | 0.72 |
| H3K4me1 | 0.87 | 0.48 | 0.56 | 0.4 | 0.74 | 0.95 | 0.85 | 1 | 0.89 | 0.76 | 0.75 | 0.61 |
| H3K4me2 | 0.8 | 0.48 | 0.67 | 0.34 | 0.71 | 0.84 | 0.79 | 0.89 | 1 | 0.94 | 0.83 | 0.65 |
| H3K4me3 | 0.68 | 0.47 | 0.7 | 0.35 | 0.66 | 0.73 | 0.76 | 0.76 | 0.94 | 1 | 0.86 | 0.65 |
| H3K9Ac | 0.73 | 0.81 | 0.79 | 0.74 | 0.78 | 0.78 | 0.89 | 0.75 | 0.83 | 0.86 | 1 | 0.9 |
| Orc | 0.7 | 0.93 | 0.81 | 0.83 | 0.78 | 0.67 | 0.72 | 0.61 | 0.65 | 0.65 | 0.9 | 1 |

**Supplemenatry data table 2.**
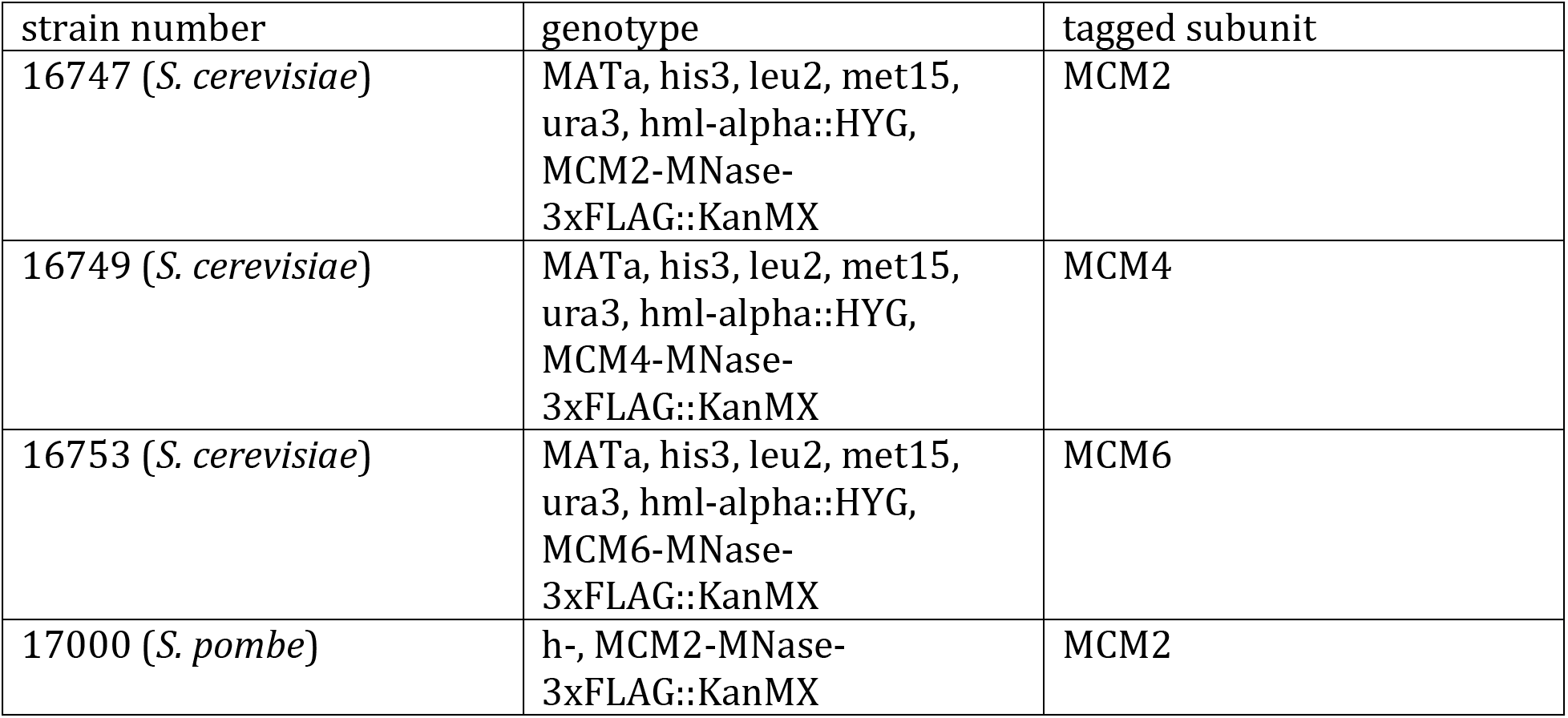
List of yeast strains

Suplemenatry data table 2. List of yeast strains
| strain number | genotype | tagged subunit |
| --- | --- | --- |
| 16747 (*S. cerevisiae*) | MATa, his3, leu2, met15, ura3, hml-alpha::HYG, MCM2-MNase-3xFLAG::KanMX | MCM2 |
| 16749 (*S. cerevisiae*) | MATa, his3, leu2, met15, ura3, hml-alpha::HYG, MCM4-MNase-3xFLAG::KanMX | MCM4 |
| 16753 (*S. cerevisiae*) | MATa, his3, leu2, met15, ura3, hml-alpha::HYG, MCM6-MNase-3xFLAG::KanMX | MCM6 |
| 17000 (*S. pombe*) | h-, MCM2-MNase-3xFLAG::KanMX | MCM2 |

